# Hiding in the yolk: a unique feature of *Legionella pneumophila* infection of zebrafish

**DOI:** 10.1101/2021.10.18.464513

**Authors:** Flávia Viana, Laurent Boucontet, Valerio Laghi, Daniel Schator, Marine Ibranosyan, Sophie Jarraud, Emma Colucci-Guyon, Carmen Buchrieser

## Abstract

The zebrafish has become a powerful model organism to study host-pathogen interactions. Here, we developed a zebrafish model of *Legionella pneumophila* infection to dissect innate immune responses. We show that *L. pneumophila* cause zebrafish larvae death in a dose dependent manner, and that macrophages are the first line of defence, with neutrophils cooperating to clear the infection. When either macrophages or neutrophils are depleted, these “immunocompromised” larvae become lethally sensitive to *L. pneumophila* similar to what is known for humans that develop pneumonia. Also as observed in human infections, the adaptor signalling molecule Myd88 is not required to control disease in the larvae. Furthermore, proinflammatory cytokine genes *il1ß* and *tnfα* were upregulated during infection, recapitulating key immune responses seen in human infection. Strikingly, we uncovered a previously undescribed infection phenotype in zebrafish larvae, whereby bloodborne, wild type *L. pneumophila* invade and grow in the larval yolk region, a phenotype not observed with a type IV secretion system deficient mutant that cannot translocate effectors into its host cell. Thus, zebrafish larva represents an innovative *L. pneumophila* infection model that that on one hand mimics important aspects of the human immune response to *L. pneumophila* infection and that on the other hand will allow to elucidate the mechanisms by which type IV secretion effectors allow *L. pneumophila* to cross membranes and to obtain nutrients from nutrient rich environments.

## INTRODUCTION

*Legionella pneumophila*, a gram negative, facultative intracellular bacterium inhabits natural, freshwater sources [1, 2]. As an environmental, aquatic microbe *L. pneumophila* replicates intracellularly in aquatic protozoa [3]. Most interestingly, in contrast to other intracellular pathogens *L. pneumophila* is not adapted to a single host, but it exhibits a broad host range including Amoebozoa (amoebae), Percolozoa (excavates) and Ciliophora (ciliated protozoa) [3, 4]. In the environment *L. pneumophila* can also be found within biofilms where it acquires nutrients from this mixed community, but it can also survive in a planktonic form for a certain time [5]. As fresh water and man-made systems are connected, *L. pneumophila* can also contaminate artificial water systems. Protected within its protozoan hosts *L. pneumophila* survives water disinfectants and may infect humans *via* aerosols produced by different man-made structures and devices. The inhalation of *L. pneumophila-contaminated* aerosols can cause a severe pneumonia, the so-called Legionnaires’ disease [6]. However, not every infection leads to disease. Disease outcome is determined by virulence of the bacterial strain, the bacterial burden in the inhaled aerosols and most importantly by the host immune status. Host factors determining susceptibility include age above 50, smoking and/or having chronic lung disease, being immunocompromised and genetic factors that alter the immune response [2, 7, 8].

Once the bacteria reach the lungs of susceptible individuals, they can infect alveolar macrophages and replicate therein. After being phagocytosed *L. pneumophila* avoids lysosomes and establishes an endoplasmic reticulum derived vacuole named the *Legionella* containing vacuole (LCV) [9, 10]. The LCV, a safe haven for bacterial replication, is established by utilizing the Dot/Icm type IV secretion system (T4SS) that injects over 330 proteins into the host cell [9–11]. These effector proteins manipulate a myriad of host pathways to recruit vesicles derived from the endoplasmic reticulum to the LCV, to supply the bacteria with nutrients, restrain autophagy and supress apoptosis and to subvert the host cell immune response [9–11]. A surprising high number of these effectors mimic host proteins and encode eukaryotic functions helping *L. pneumophila* to modulate numerous host pathways to its own benefit in remarkable diverse ways [11–13]

Intracellular replication of *L. pneumophila* and innate immune responses to this pathogen have been studied *in vitro* using both murine and human cell lines and *in vivo* using different animal models of infection. However, results obtained with these models cannot be easily extrapolated to what is observed in human disease. Studies in invertebrate models, such as *Galleria mellonella* and *Caenorhabditis elegans*, [14, 15] require further validation in more developed models as their immune system greatly differs from that of vertebrates. Interestingly, mouse infection fails to recall the human disease phenotype, as most inbred mice strains are naturally resistant to *L. pneumophila* [16]. Very early after the discovery of *L. pneumophila* the guinea pig model of Legionnaires’ disease was developed as these are highly susceptible to *L. pneumophila* when infected through injection into the peritoneum [6] or when exposed to *L. pneumophila* containing aerosols [6]. Several studies thereafter have shown that the guinea pig infection model recalls human disease and allows to study the immune response to *L. pneumophila* infection [17, 18]. However, the guinea pig model is rarely used due to the limited availability of specific immunological reagents for these animals and the demanding laboratory and husbandry requirements.

The above-mentioned models, including the widely used murine models, present limitations for studying *L. pneumophila* infection *in vivo* and discrepancies exist between results obtained in mouse or human cells, as for example mouse macrophages restrict *L. pneumophila* growth via caspase 1 and caspase 7 activation whereas human macrophages do not activate caspase 1 and 7 and thus allow growth of *L. pneumophila* [19, 20]. Thus, we sought to develop a new, alternative model for *Legionella* infection. The zebrafish (*Danio rerio*) originally introduced as a model organism in developmental biology has emerged in recent years as a powerful non-mammalian model to study nearly every aspect of biology, including immune cell behaviour and host-pathogen interactions [21, 22]. Zebrafish are evolutionary closer to humans than fruit flies and nematodes, easier to manipulate than mice and their immune system is remarkably similar to the one of mammals, making them an attractive laboratory model for immunology and infection biology [21, 22]. Its popularity is also due to its small size and the natural translucency of its embryos and larvae, which makes it possible to follow leukocyte behaviour and infection onset at the level of the whole organism in real-time and high resolution [23]. Additionally, although adult organisms display a fully developed immune system with both active innate and adaptive branches, studies can also be conducted at the early stages of life (embryonic or larvae) when the organism solely relies on innate immunity, allowing to dissect the mechanisms arising from different immune responses [23–25]. Here we examined whether the zebrafish could be an alternative model for analysing host-pathogen interactions and the innate immune response to *L. pneumophila* infection.

We show that *L. pneumophila* infection of zebrafish larvae recapitulate human disease onset, as infected wild-type larvae are generally able to clear the infection, but immunocompromised fish fail to do so. Both macrophages and neutrophils quickly interact and engulf injected *L. pneumophila*. Macrophage-depleted larvae show a dramatic increase of bacterial burden concomitant with host death, pointing to a crucial role of macrophages in controlling the infection. However, not all larvae are able to control the infection, but a fraction showed high bacterial burden in the yolk region, a unique zebrafish infection phenotype.

## RESULTS

### L. pneumophila infection induces mortality in zebrafish larvae in a dose dependent manner

To analyse whether *L. pneumophila* can cause disease in zebrafish larvae we microinjected larvae 72 hours post fertilisation (hpf) intravenously in the caudal vessels near the cloaca (Fig 1A), with 10^3^ or 10^4^ CFU of wild type (WT) *L. pneumophila* strain Paris expressing GFP (WT-GFP) or the type IV secretion system (T4SS) deficient isogenic mutant expressing GFP (*ΔdotA-GFP*). The infected larvae were kept at 28°C and were monitored regularly until 72 hours post infection (hpi) to record survival or death using a stereomicroscope. Larvae infected with doses of up to 2×10^3^ CFU of WT-GFP (defined as low dose, LD) all survived (100% survival). In contrast, larvae infected intravenously with doses of 10^4^ CFU (defined as high dose, HD) resulted in approximately 30% of death within 72 hpi (Fig 1B). Importantly, all larvae injected with HD of the Δ*dotA*-GFP strain survived for the entire time of observation (Fig 1B) indicating that the T4SS is crucial for replication in zebrafish larvae as it is in other infection models and in humans.

**Figure 1.**
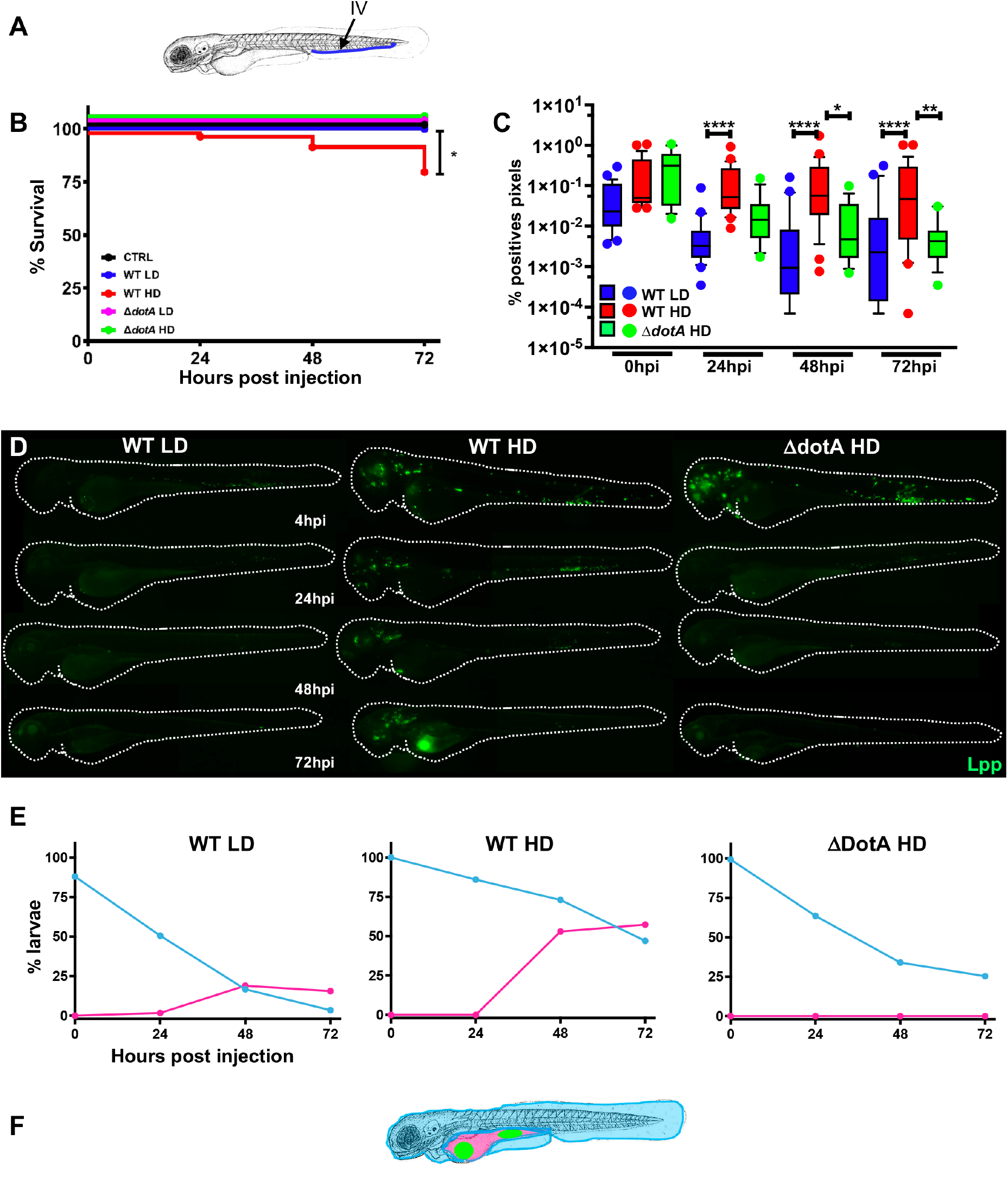
Zebrafish larvae are susceptible to intravenous *L. pneumophila* infection in a dose dependent manner. **A)** Scheme of the experimental set up of bacterial infection using zebrafish. A 72hpf zebrafish larva is shown. Bacteria are injected in the bloodstream (iv) via the caudal vein (green arrow). **B)** Survival curves (three independent experiments pooled) of zebrafish larvae injected with WT-GFP Low Dose (WT LD) (blue curve, n=60) or High Dose (HD) (red curve, n=60), or with Δ*dotA*-GFP Low Dose (Δ*dotA* LD) (green curve, n=12) or High Dose (Δ*dotA* HD) (green curve, n=36), and incubated at 28°C. Control non-injected fish (CTRL, black curve; n= 72). P < 0.05 was considered statistically significant (symbols: **** P < 0.0001; ***P < 0.001; **P < 0.01; *P < 0.05. **C**) Bacterial burden evaluation by quantification of % of fluorescent pixel counts from images of individual injected larvae followed overtime from 0 to 72hpi. Each larva was imaged daily, and images were analysed with Fiji for bacterial burden quantification. Five experiments have been pooled, for a total of 28 larvae for WT LD and LD, 18 for Δ*dotA* HD. P < 0.05 was considered statistically significant (symbols: **** P < 0.0001; ***P < 0.001; **P < 0.01; *P < 0.05. No symbol on graphs means that not statistically differences were observed. -**D)** Representative images of *L. pneumophila* dissemination, determined by live imaging using a fluorescence stereomicroscope, of zebrafish AB larvae infected with a LD or a HD of WT-GFP, or a HD of Δ*dotA*-GFP. Individual infected larvae were followed over time by live imaging at 4h, 24h, 48h, and 72h post injection. GFP fluorescence of the injected bacteria is shown. **E)** Quantification of bacterial dissemination overtime. Larvae injected with LD, HD LD or Δ*dotA* HD were imaged overtime, and then scored for the GFP + bacteria absolute presence for each larva overtime. Larvae were scored as “infected” when they showed at least one small detectable GFP+ dot. Data obtained were plotted as % of larvae with bacteria in the body (tail, trunk, head; blue curve), or in the yolk (pink curve). 11 independent experiments have been plotted (representing a total of n=69 WT LD; n=58 WT HD; n=54 Δ*dotA* HD infected larvae). **F**) Scheme of a 72hpi larva with body (light blue) and yolk (pink) region highlighted. Green dot in the yolk represents bacteria burden.

We then set up a method to monitor the bacterial burden of the infected zebrafish larvae. The progression of the infection was followed by analysing the bacterial load at 0, 24, 48 and 72 hpi comparing three different methods. First, we quantified the pixel counts of GFP fluorescence of live larvae images (Fig. S1A), secondly, we analysed the number of GFP expressing bacteria present in lysed infected larvae by FACS (Fig. S1B) and thirdly we plated serial dilutions of homogenates of euthanized larvae on BCYE medium (Fig S1C). The results obtained with the three methods showed a similar trend, but the method using fluorescent pixel counts on live injected larvae seemed to most accurately represent the bacterial load. Thus, we choose to monitor the *L. pneumophila* load of zebrafish larvae by fluorescent pixel counts on live injected larvae, and in addition by FACS on lysed larvae at various time points, using the GFP fluorescence of the bacteria. As shown in Fig. 1C, where the bacterial burden was evaluated by fluorescent pixel counts on individual injected larvae followed over 72hpi, most larvae injected with LD of WT-GFP progressively control the bacteria by 24 hpi with only few larvae showing an increase in bacterial numbers at 72hpi. Similarly, HD of Δ*dotA*-GFP were also progressively controlled by 24 hpi. In contrast, some zebrafish larvae injected with HD of WT-GFP were unable to eliminate the bacteria at 72hpi, and the bacterial counts remained high and increased by 48-72 hpi (Fig 1C). These results were corroborated by FACS analysis on lysed larvae (Fig S2 A). We also monitored infected larvae by fluorescence microscopy. Immediately upon injection (20 min to 2 hpi), bacteria were detectable as small foci, probably associated with professional phagocytes (Fig. 1D). By 24 hpi, in both, larvae injected with LD of WT-GFP as well as larvae injected with HD of the avirulent Δ*dotA*-GFP strain, the GFP signal declined becoming undetectable by 48 hpi, suggesting that the bacteria were progressively cleared. Despite showing the same pattern at 24 hpi, larvae injected with HD of WT-GFP showed an increase in GFP signal at 48 hpi, suggesting that bacterial proliferation occurred in a fraction of the infected larvae. Interestingly, in these larvae, bacterial proliferation occurred mainly in the yolk region where the bacterial foci increased dramatically over time, with concomitant death of the infected larvae by 72 hpi (Fig 1D).

To gain insight into the progression of the infection, we analysed the bacterial presence in the yolk or in the rest of the body (tail + trunk + head) of LD and HD WT-GFP or HD Δ*dotA*-GFP infected larvae under the fluorescence microscope over time. A single small GFP dot (indicating few bacteria) present in the larvae was scored as “positive for infection”. We observed that about 60% of larvae injected with HD WT-GFP and about 20% of larvae injected with LD WT-GFP showed yolk growth at 72 hpi. In contrast, larvae injected with HD Δ*dotA*-GFP progressively cleared the bacteria, and bacteria were never observed growing in the yolk (Fig 1E, F; Figure 3D). The presence of bacteria in the yolk was intriguing and prompted us to investigate whether this unique feature was dependent on the site of injection of bacteria in the larvae. Thus, we injected 72hpf zebrafish larvae with HD WT-GFP in different closed cavities such as the otic vesicle and the hind brain ventricle. The survival, bacterial burden, and outcome of bacterial dissemination over time compared to bloodstream injected bacteria showed that only blood borne bacteria can successfully replicate in zebrafish and establish a proliferative niche in the yolk. This suggests a role of the blood circulation in the capacity of *L. pneumophila* to reach the yolk region and to replicate there (Fig S3 A-C).

**Figure 2.**
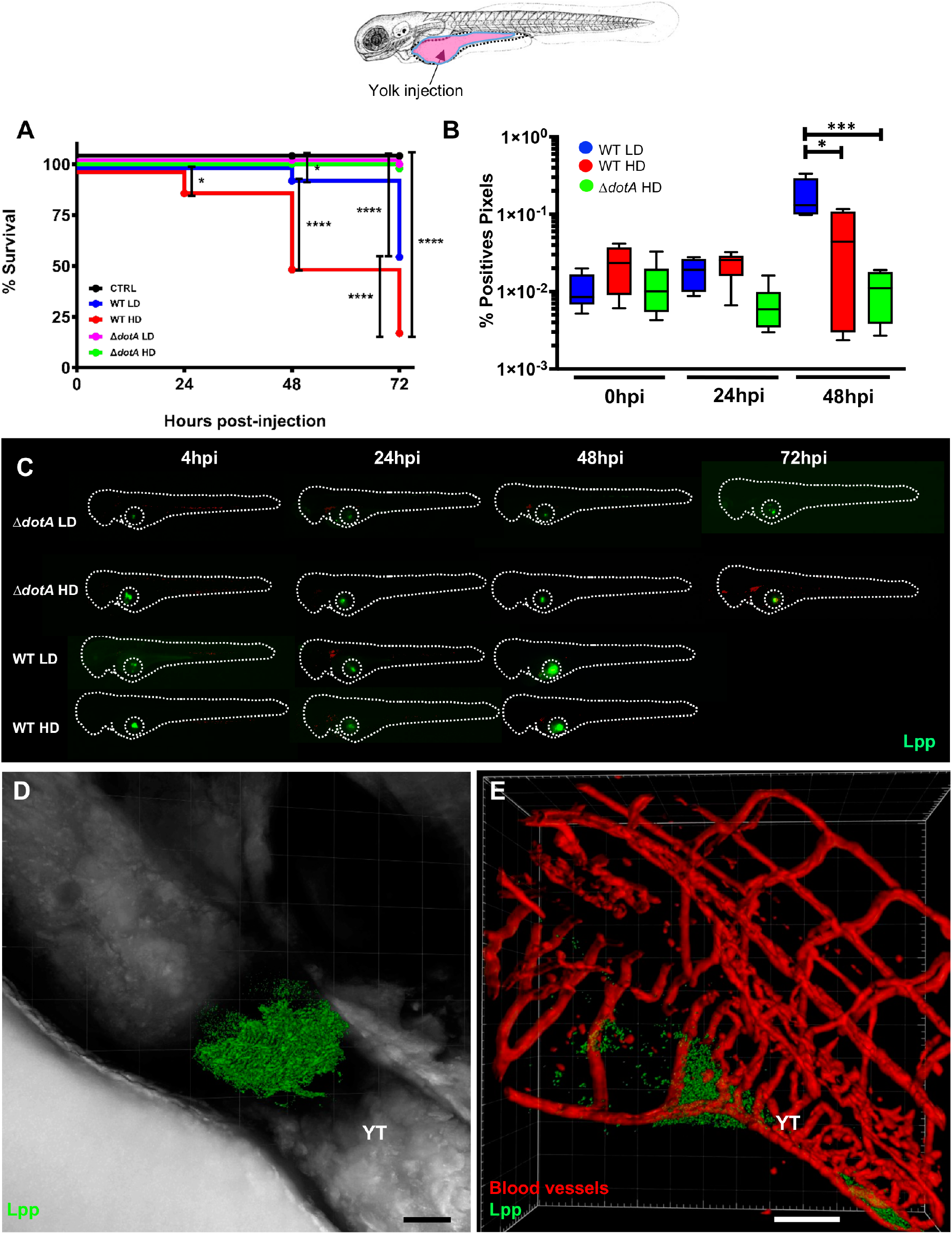
Bloodstream *L. pneumophila* establish a proliferative niche in the yolk causing a persistent local infection. **A)** Survival curves (two independent experiments pooled) of zebrafish larvae injected with WT-GFP Low Dose (WT LD) (blue curve, n=48) or High Dose (HD) (red curve, n=48), or with Δ*dotA*-GFP Low Dose (Δ*dotA* LD) (magenta curve, n=48) or High Dose (Δ*dotA* HD) (green curve, n=48), and incubated at 28°C. Non-injected fish (CTRL, black curve; n= 48). **B)** Bacterial burden evaluation by quantification of % of fluorescent pixel counts on individual injected larvae followed overtime by 0 to 48hpi. Each larva was imaged daily, and images were analysed with Fiji for bacterial burden evaluation. One experiment plotted, 6 larvae per condition. P < 0.05 was considered statistically significant (symbols: **** P < 0.0001; ***P < 0.001; **P < 0.01; *P < 0.05. No symbol on graphs means that not statistically differences were observed. **C)** Representative images of *L. pneumophila* localization, determined by live imaging using a fluorescence stereomicroscope, of zebrafish larvae infected with a LD or a HD of WT-GFP, or a LD or a HD of Δ*dotA*-GFP. Individual infected larvae were followed over time by live imaging at 4h, 24h, 48h, and 72hpi. GFP fluorescence of the injected bacteria is shown. Dotted circle highlights GFP bacteria in the yolk. **D)** Representative maximum intensity projection of confocal acquisition of a 72hpi zebrafish larva injected in the bloodstream with HD of WT-GFP, mounted laterally and live imaged using high resolution confocal fluorescence microscope, showing bacteria growing in the yolk and yolk tube region. The x-y-z raw data were post treated with the LEICA lighting application for reducing noise and processed with Imaris for 3D volume rendering. Related to Movie S1. **E)** Imaris 3D reconstruction and volume rendering of the *L. pneumophila* growth (GFP labelling) in the yolk of *kdrl*:ras-mCherry (red vessels) infected larva at 72hpi, showed laterally. Overlay of GFP and mCherry is shown; BF is shown to help to visualize the yolk region and host anatomy. Related to Movie S2. Scale bar = 50mm.

**Figure 3.**
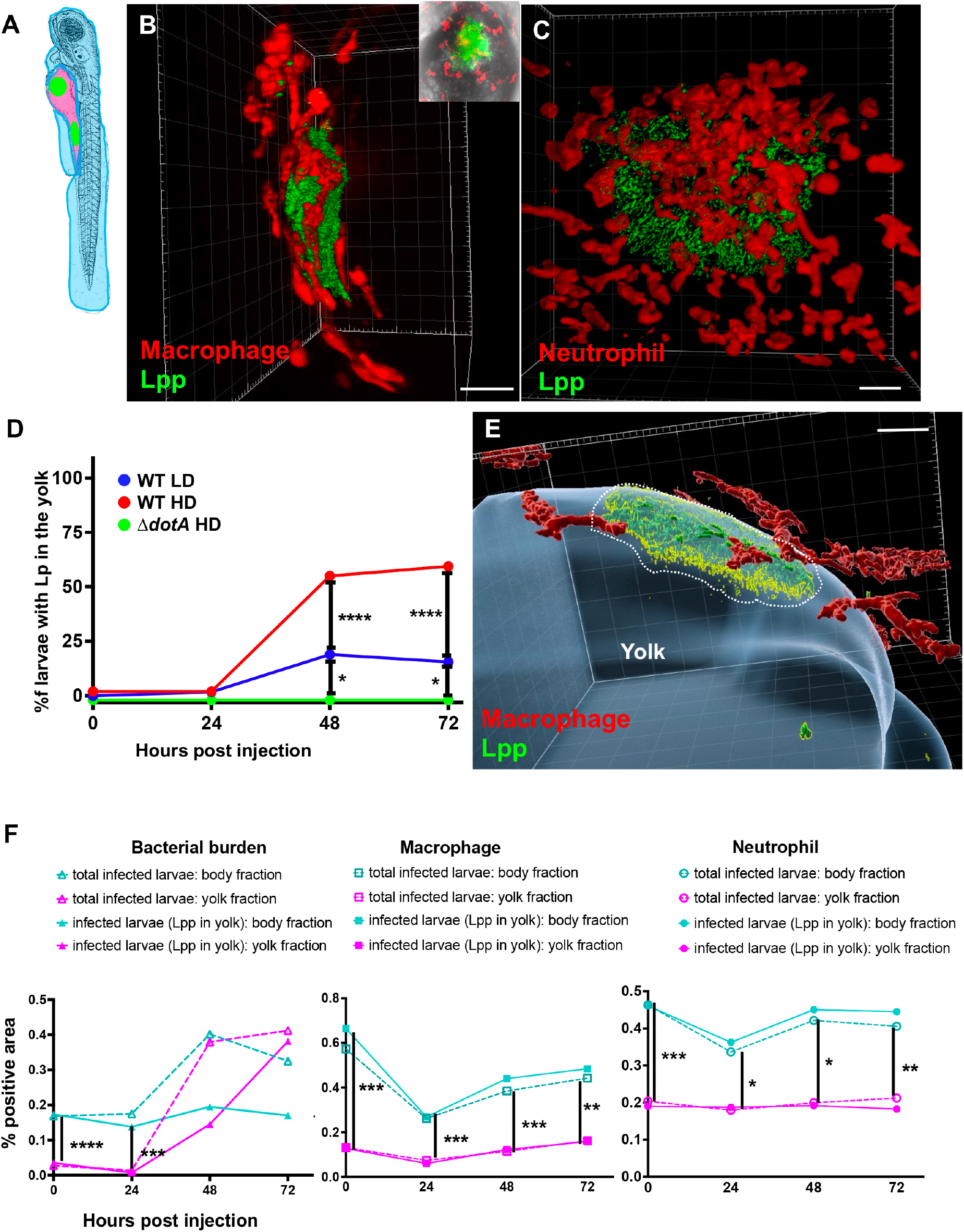
Characterization of the *L. pneumophila* foci growing in the yolk region of zebrafish larvae. **A)** Scheme of 72hpf with body (light blue) and yolk region (pink) highlighted; the yolk sustaining *L. pneumophila* growing has been indicated with green dots**. B**) Imaris 3D reconstruction and volume rendering of the *L. pneumophila* growth in the yolk of 72hpi *mfap4:* mCherry larva (red macrophages) injected in the bloodstream with HD of WT-GFP at 72hpf, shown laterally. Inset shows the maximum intensity projection of the *L. pneumophila* foci of the same larva mounted ventrally. Scale bar = 20mm. Related to Movie S3. **C)** Imaris 3D reconstruction and volume rendering of the *L. pneumophila* growth in the yolk of *lyz*:DsRed (red neutrophils) infected larva at 72hpi, showed laterally. Scale bar = 20mm. Related to Movie S4. **D)** Quantification of bacterial burden in the yolk overtime. Larvae injected with LD, HD WT, or Δ*dotA* HD were imaged overtime, and then scored for the GFP + bacteria absolute presence in the yolk for each larva overtime. Larvae were scored as “infected” when they showed at least one small detectable GFP+ dot. Data obtained were plotted as % of larvae with bacteria in the in the yolk upon LD (blue curve), HD (red curve) WT or Δ*dotA* HD (green curve) injection overtime. 11 independent experiments have been plotted (representing a total of n=69 WT LD; n=58 WT HD; n=54 Δ*dotA* HD infected larvae) **E)** Time frame extracted from a 4D acquisition of *L. pneumophila* growth in the yolk between 48 and 72hpi of *mfap4:* mCherry larva (red macrophages) injected in the bloodstream with HD of WT-GFP shown laterally. Imaris 4D reconstruction and volume rendering of the bacteria aggregate and interaction with macrophages. The yolk has been manually highlighted (in blue) with Imaris. Scale bar: 10. Related to Movie S5. **F)** Quantification of bacterial burden in the whole body, in the body or in the yolk region versus macrophage or neutrophil quantification in the body or in the yolk region in HD WT-GFP infected larvae followed overtime. Two independent experiments plotted for each phagocyte type (total of 11 larvae for macrophage and 11 larvae for neutrophil quantification). Quantification of the fluorescence images (GFP bacteria and RFP leukocytes) was done using CellProfiler software. P < 0.05 was considered statistically significant (symbols: **** P < 0.0001; ***P < 0.001; **P < 0.01; *P < 0.05). No symbol on graphs means that not statistically differences were observed.

Secondly, we tested if a natural route of uptake could cause infection in zebrafish. As in its usual habitat *L. pneumophila* lives in freshwater and replicates in protozoan hosts [26] it is possible that fish get infected in the environment by taking up infected amoeba. Indeed, amoebae are prey of zebrafish larvae. To test this hypothesis, we infected *A. castellanii* with *L pneumophila* and exposed 120hpf zebrafish larvae (start of autonomous feeding) to *L. pneumophila* infected amoebae and evaluated bacterial survival (Fig S4 A) and bacterial dissemination (Fig S4 B-D) in the larvae over time. While zebrafish larvae engulfed the infected amoebae, as shown by the GFP signal detectable in the exposed larvae at 48 hours post contact bath (Fig S4C), no permanent infection was established and the engulfed bacteria were evacuated with other faecal content without impact on larvae survival, suggesting that this might not be an important route of infection in the environment.

Collectively these results indicate that only bloodstream injected WT *L. pneumophila* induce a dose dependent death of zebrafish larvae. Larvae that were unable to control infection by 48 hpi, showed a unique infection phenotype, a high increase of the bacterial burden in the yolk region.

### Legionella pneumophila replication in the yolk of zebrafish larvae is T4SS dependent

The replication of *L. pneumophila* in the yolk region of infected zebrafish larvae was strictly dependent on a functioning T4SS as Δ*dotA-*GFP failed to reach the yolk. To investigate whether the secretion mutant can grow in the yolk cell when reaching it, we injected LD and HD of WT-GFP or of Δ*dotA-*GFP *L. pneumophila* directly into the yolk cell cytoplasm of 72hpf zebrafish larvae (Fig. 2A). Surprisingly, Δ*dotA*-GFP did not replicate in the yolk even when injected directly although it persisted over 48hpi (Fig 2B, C, Fig S2B). WT-GFP replicated in the yolk region both, upon LD and HD injections resulting in an increased bacterial proliferation in the yolk and subsequent death of the larvae whereby 100% of the larvae dying have bacteria replicating in the yolk. (Fig 2B, C, Fig S2B). These observations suggest that T4SS system is not only crucial for reaching the yolk region but likely that some of its effectors are necessary for the bacteria to obtain nutrients from the yolk environment to allow replication. To further analyse this hypothesis, we selected a mutant in the gene encoding a sphingosine-1 phosphate lyase, (WT, Δ*spl*) [27] as we reasoned that this enzyme might be implicated in degrading sphingolipids present in the yolk of zebrafish larvae and thereby might aid *L. pneumophila* to obtain nutrients. Injection of Δ*spl* in the yolk sac region, and analyses of larvae death as compared to WT or Δ*dotA* showed that survival of zebrafish larvae injected with the Δ*spl* was slightly but significantly higher than with WT injected larvae (Fig. S5A), suggesting that the T4SS effector *Lp*Spl might be implicated in nutrient acquisition in the yolk environment.

Interestingly, the first isolation of *L. pneumophila* was achieved by inoculating the yolk region of embryonated eggs probably due to the richness in nutrients provided by the yolk [6]. Thus, we decided to investigate the infection phenotype of *L. pneumophila* WT and Δ*dotA* in the yolk sac further using as model embryonated chicken eggs (ECE). We inoculated ECE directly in the yolk region with WT and with the Δ*dotA* strain at a concentration of 9.2 log_10_ CFU/mL and 9.1 log_10_ CFU/mL, respectively and assessed mortality of the embryos daily. The survival curves were significantly different (p=0.0111). The total mortality during the 6-day observation period was significantly higher in WT-GFP infected eggs (88.9%) than in the Δ*dotA-*infected eggs (14.3%) or PBS inoculated control eggs (28.6%) (Fig. S5C). The highest mortality was observed at 2 days post infection in WT inoculated eggs with 55.6% mortality *versus* 0% in Δ*dotA* or 28.6% mortality in PBS inoculated eggs. Quantification of *L. pneumophila* in the yolk sac region at the day of mortality or at day 6 post infection revealed that the number of bacteria in the yolk sac of WT-infected ECE, was significantly higher than that in the yolk sac of those infected with the Δ*dotA* strain (7.8 log_10_ CFU/mL and 5.9 log_10_ CFU/mL, respectively, p=0.0127) (Fig. S5D). Controls inoculated with PBS (n=2) showed no *L. pneumophila* growth. Thus, like in zebrafish larvae only the WT strain can replicate in the yolk region and of inducing mortality in the embryos, while the T4SS mutant strain persists but is not able to replicate and does not induce high embryo mortality. This result further supports the finding that the T4SS system is crucial for obtaining nutrients when lipids are the major available energy source.

Taken together, these results suggest that the *L. pneumophila* T4SS plays a crucial role for the bacteria to pass from the blood circulation into the yolk and that T4SS effectors play an important role to obtain nutrients for bacterial proliferation in the yolk.

### Bloodstream L. pneumophila establishes a proliferative niche in the yolk region causing a persistent infection

To characterise the *L. pneumophila* foci identified in the yolk region of zebrafish larvae, we used high resolution fluorescence microscopy of HD of WT-GFP *L. pneumophila* injected in the bloodstream of 72hpf zebrafish larvae and analysed them at 72hpi. This analysis confirmed that these bacterial structures localize in the yolk and or in the yolk tube region (Fig. 2D, Movie S1). *L. pneumophila* foci in the yolk region are highly complex, aggregate-like structures of long, filamentous bacteria. Moreover, upon injection of HD WT-GFP in Tg(*kdrl*::mCherry)^is5^ (red blood vessels) larvae, we showed that the fast growing bacterial aggregates localize close to the blood vessels, mostly below and probably above the yolk cell. Single bacteria near the aggregates localized within the blood vessels, indicating that the bacteria can cross the endothelial barrier to reach the yolk region (Fig 2E, Movie S2). To analyse macrophage and neutrophil interactions with the bacterial aggregates in the yolk region, we injected HD WT-GFP in Tg(*mfap4::mCherryF*) (herein referred as *mfap4*:mCherryF) (red macrophages) or Tg(*Lyz::DsRed)*^nz50^ (herein referred as *lyz*:DsRed)(red neutrophils) or Tg(*kdrl*::mCherry)^is5^ (red blood vessels) zebrafish larvae. At 72hpi macrophages accumulated around the yolk region containing *L. pneumophila* but did not seem to be able to engulf the bacterial aggregates. (Fig. 3A, Movie S3). Similarly, upon injection of HD of WT-GFP in *lyz*:DsRed larvae (red neutrophils), at 72hpi neutrophils accumulated around the growing bacterial aggregates, but seemed also unable to engulf them (Fig 3B, C, Movie S4). Strikingly, quantification analyses showed that bacterial colonisation of the yolk of zebrafish larvae injected with the T4SS deficient Δ*dotA* mutant strain, did never take place, suggesting that zebrafish susceptibility to *L. pneumophila* infection and yolk penetration depends on a functional T4SS system (Fig. 3D). It should be noted that the yolk is the only food source of the larvae during this developmental stage. The fast proliferation of the bacteria in the yolk region probably depletes its nutritional content, leading to larvae death.

To gain deeper insight into the exact anatomical localisation of the bacteria in the yolk region, we performed high resolution confocal time lapse acquisitions of HD of WT-GFP bloodstream injected larvae harbouring red macrophages between 48 and 72hpi. We observed that *L. pneumophila* foci seem to localize both above but also below the plasmatic yolk membrane, suggesting that they can cross it, and that professional phagocytes remain above the growing bacterial aggregates failing to engulf them (Fig 3E, Movie S5). Live imaging showed that both macrophages and neutrophils accumulated around bacteria proliferating in the yolk region. To investigate if professional phagocytes were recruited to the bacteria from other sites, or if only the population located on the yolk was involved, we quantified macrophages and neutrophils in the body and in the yolk of the whole larva overtime, focusing on HD WT-GFP infected larvae. We then separated the larvae with bacterial burden in the yolk from the total population of infected larvae, to specifically analyse the recruitment of professional leukocytes to the bacteria growing in the yolk. Following the same criteria as above, we quantified the bacterial burden of the infected larvae over time. This quantitative analysis showed that macrophages and neutrophils that accumulated where the bacteria are seen in the yolk by 48hpi, were the ones located on the yolk surface, and that no professional phagocyte population was recruited in the other parts of the body (Fig 3F; Fig S6). Moreover, this analysis confirmed that the bacterial burden increased in the yolk while decreasing in the body (Fig 1E) and it also revealed a decrease of professional phagocyte populations in infected larvae upon HD WT-GFP injection overtime.

Thus, blood-borne *L. pneumophila* can invade the yolk sac of zebrafish larvae, a previously undescribed phenotype of bacterial infection in this model. Once in the yolk, the bacteria replicate extensively, forming complex, organized, aggregate-like structures that cannot be removed by macrophages and neutrophils, thereby avoiding the host’s immune control and clearance, and eventually causing the death of the larvae.

### Infection of zebrafish larvae with high doses of L. pneumophila leads to macrophage and neutrophil death

In human infection, alveolar macrophages are the primary cell type infected by *L. pneumophila* supporting its intracellular replication. Following infection, neutrophils are recruited to the lung and are key players for controlling infection as they possess antimicrobial activity and kill *L. pneumophila* [28]. Moreover, the analysis of the recruitment of professional leukocytes to the bacteria in the yolk of HD WT-GFP infected larvae revealed a decrease of professional phagocyte populations overtime (Fig 3F). Thus, to analyse whether zebrafish infection mirrors human infection, we monitored the behaviour of zebrafish macrophages or neutrophils over time. The transgenic zebrafish larvae *mfap4:mCherryF* and *lyz*:DsRed were injected with low or high doses of WT-GFP or with high doses of Δ*dotA*-GFP. Infected live larvae were monitored using widefield fluorescence microscopy and the number of leukocytes per larva was assessed by counting fluorescent macrophages and neutrophils over time until 72hpi. We observed that upon injection of high dose WT-GFP, the macrophage count decreased dramatically at 24hpi and 48hpi, but started to increase at 72hpi (Fig. 4A). Neutrophil counts gave similar results at 24 and 48hpi upon injection of high doses of WT bacteria, as there was a dramatic decrease observed in neutrophil numbers by 24hpi. However, the neutrophil counts were still decreased at 72hpi (Fig. 5A). In contrast macrophage and neutrophil counts remained unaffected upon injection of equal amounts of the avirulent Δ*dotA* strain, with a slight increase of neutrophil numbers at 72hpi, suggesting that phagocyte death is linked to a functional T4SS system (Fig 4A, 5A).

**Figure 4.**
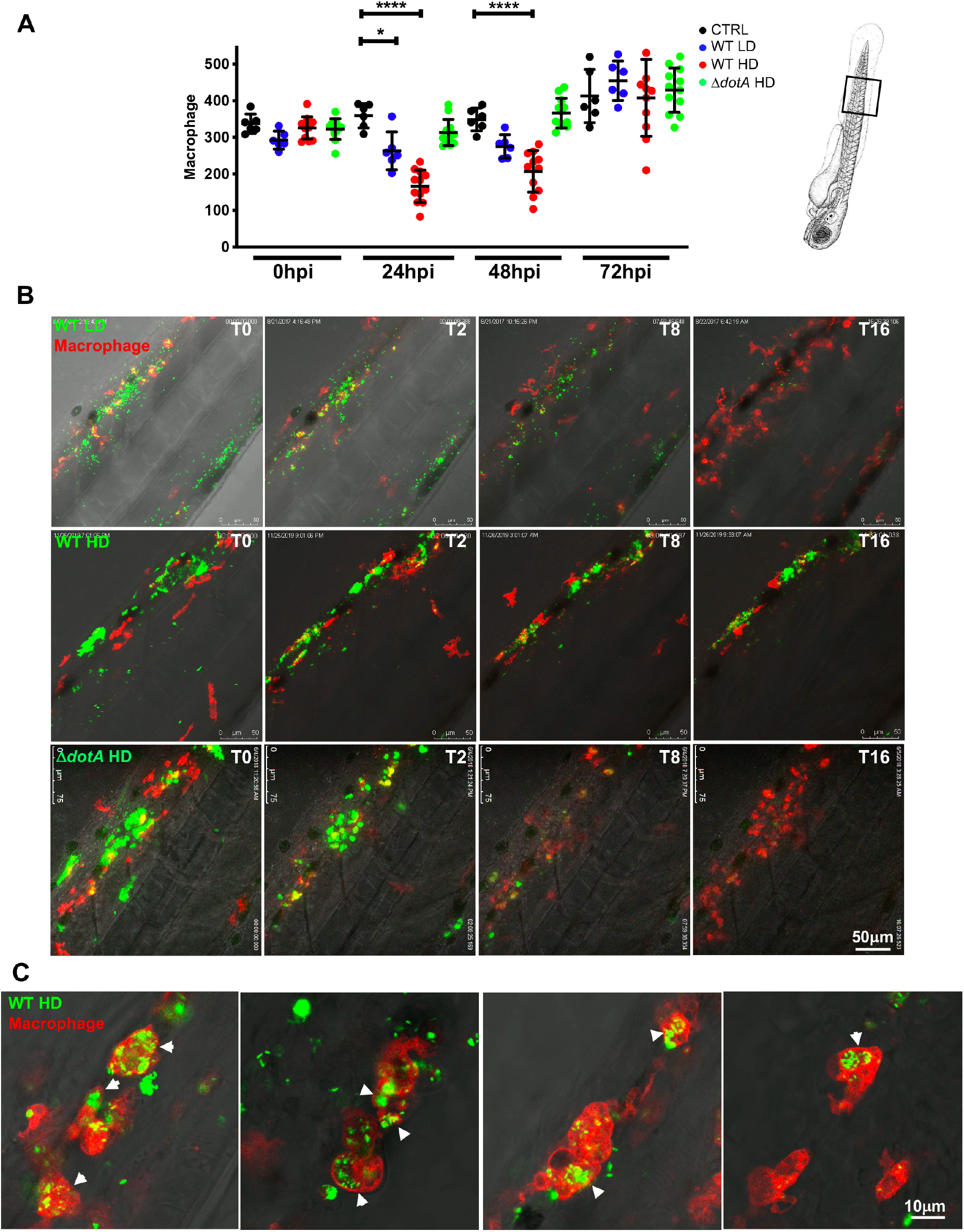
*L. pneumophila* high dose injection results in (systemic) macrophage and neutrophil death. **A)** Macrophage counts of control larvae (black symbols) or upon Low Dose (blue symbols) or High Dose of WT-GFP (red symbols), or High Dose (green symbols) of Δ*dotA* -GFP injection. Macrophages were counted manually from images taken on live infected larvae over time from T0 to T72hpi, using ImageJ software, and results were plotted using GraphPad Prism^®^ software. Mean±SEM are also shown (horizontal bars). Data plotted are from two pooled independent experiments (n=12 larvae scored for each condition). P < 0.05 was considered statistically significant (symbols: **** P < 0.0001; ***P < 0.001; **P < 0.01; *P < 0.05). No symbol on graphs means that not statistically differences were observed. **B)** Frames extracted from maximum intensity projection of *in vivo* time-lapse confocal fluorescence microscopy of 72hpf Tg(*mfap4::mCherryF*) larvae injected in the bloodstream (iv) with a LD, HD (of WT-GFP or a HD of Δ*dotA*-GFP (upper panel) or Tg(*LysC::DsRed)^nz50^* in the bloodstream with a LD, HD of WT-GFP or a HD of Δ*dotA*-GFP (lower panel) to follow macrophage bacteria interaction overtime during the first 16hpi. Overlay of green (*L. pneumophila*) and red (leucocytes) fluorescence of the caudal area of the larvae (region boxed in the scheme on the right of the panel) is shown. BF helps for anatomical region indication. Representative of n= 12 to 16 injected larvae for each condition. Scale bar: 50⍰m. See also related Movies S6. **C)** macrophage *L pneumophila* interaction at 72hpi captured at high resolution upon HD WT injection. Live bacteria are inside zebrafish macrophages suggesting the establishment of a replicative niche, as documented for cultured mammalian macrophages or amoebae. Representative of n= 15 scored infected larvae. Overlay of green (GFP bacteria), red (mCherry macrophages) and BF is shown. Scale bar= 10mm

**Figure 5.**
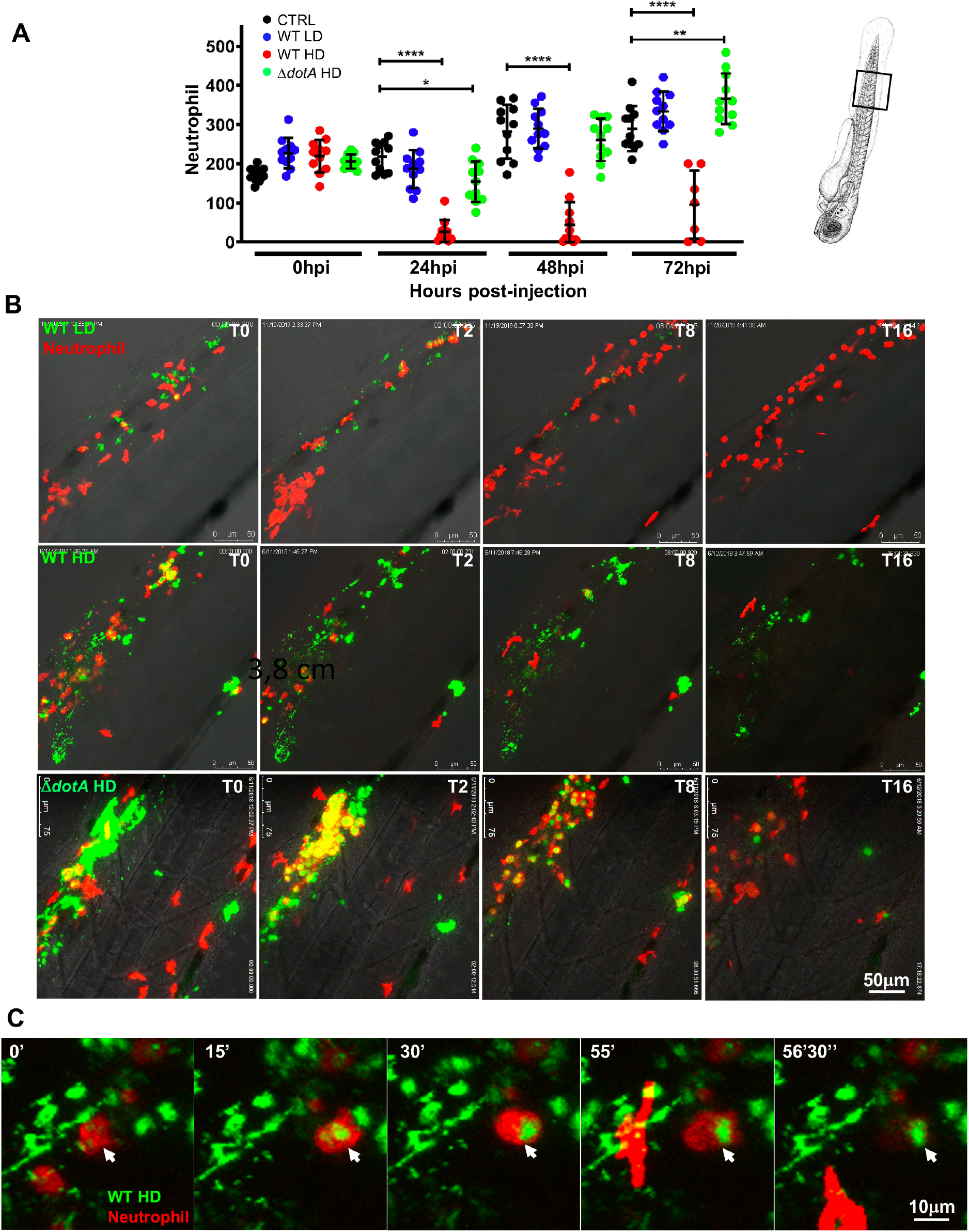
*L. pneumophila* interaction with neutrophils and neutrophil counts upon bloodstream injection of LD, HD WT GFP or HD ΔdotA *L pneumophila*. **A)** Neutrophil counts from control larvae (CTRL, black symbols) or Low Dose or High Dose of WT-GFP (blue or red symbols), or High Dose of Δ*dotA*-GFP (green symbols) injected larvae. Data plotted in the same way as for macrophage counts in Fig 4. Two pooled independent experiments (n=10 larvae scored for each condition). P < 0.05 was considered statistically significant (symbols: **** P < 0.0001; ***P < 0.001; **P < 0.01; *P < 0.05). No symbol on graphs means that not statistically differences were observed. **B)** Frames extracted from maximum intensity projection of *in vivo* time-lapse confocal fluorescence microscopy of 72hpf Tg(*LysC::DsRed)^nz50^* in the bloodstream (iv) with a LD, HD of WT-GFP or a HD of Δ*dotA*-GFP (lower panel) to follow neutrophil interaction with *L. pneumophila* immediately upon injection for 16hpi. Images were taken from time lapse at different time points (0hpi, 2hpi, 4hpi, 8hpi and 16hpi). Overlay of green (*L. pneumophila*) and red (neutrophils) fluorescence of the caudal area of the larvae (region boxed in the scheme on the right of the panel) is shown. Data representative of n= 12 to 16 larvae scored. Scale bar: 50⍰m. See also related Movies S7. **C)** Details of a dying phagocytosing neutrophil, progressively rounding-up and loosing fluorescence upon HD WT injection. Arrowheads point to the dying phagocytosing neutrophils. Representatives of n= 14 larvae scored. Scale bar=10mm. See also related Movie S8.

Taken together, these results show that high dose *L. pneumophila* infection leads to a decrease in the number of professional phagocytes dependent on the T4SS, like what is seen during human infection by *L. pneumophila* or *Mycobacterium tuberculosis* [28, 29]

### Macrophages are the primary cells to phagocytise blood-borne L. pneumophila and neutrophils cooperate to decrease bacterial load

As macrophages and neutrophils are the phagocytes known to firstly interact with *L. pneumophila* we analysed phagocyte-*L. pneumophila* interactions *in vivo* by injecting *mfap4:mCherryF* or *lyz*:DsRed 72hpf larvae with WT-GFP or Δ*dotA*-GFP and recorded phagocyte-*L. pneumophila* interactions using high resolution confocal microscopy. This showed that upon injection of LD WT-GFP, macrophages immediately contacted and engulfed blood-borne bacteria. Macrophages were continuously recruited to the site of injection and by 16hpi the bacteria were mostly undetectable while macrophage numbers increased (Fig. 4B top panel, Movie S6). Macrophages that had engulfed a large amount of *L. pneumophila* stopped moving and rounded-up. Similarly, the inhibition of the migration of phagocytes by *L. pneumophila* has been observed previously during infection of RAW 264.7 macrophages and the amoeba *Dictyostelium discoideum* and *Acanthamoeba castellanii*, [30, 31]. In contrast, zebrafish injected with HD of WT-GFP were not able to restrict the bacterial growth by 16hpi. Injection of HD of *L pneumophila* led to the formation of big bacterial aggregates, that were not easily engulfed and cleared by macrophages, as previously shown for bacterial aggregates proliferating in the yolk region (Fig 4B, second panel, Movie S6). Remarkably, macrophages were very efficient in engulfing and rapidly clearing high doses and big bacterial aggregates of blood-borne Δ*dotA*-GFP bacteria. By 10hpi most of the Δ*dotA* bacteria and bacterial aggregates had been engulfed and cleared as suggested by the diffuse GFP staining in phagocytes (Fig. 4B, bottom panel, Movie S6). However, upon infection with a HD WT-GFP, bacteria were not completely cleared and at 72hpi live *L. pneumophila* were found in macrophages, suggesting that the bacteria are also replicating in macrophages of zebrafish larvae. Indeed, high resolution confocal microscopy showed that at 72hpi, *L. pneumophila* can also be found inside of macrophages in structures resembling replicative vacuoles (Fig. 4C).

The analyses of *L. pneumophila-neutrophil* interactions showed that these cells engulfed the bacteria trapped in the mesenchyme around the site of injection, but they were less efficient at clearing blood-borne bacteria. This has also been previously observed for infection of zebrafish larvae with *Escherichia coli* or *Shigella flexneri* [24, 32]. Indeed, upon injection with a HD of WT-GFP, neutrophil failed to restrict *L. pneumophila*, leading to massive death of infected neutrophils, they rounded up and lost their fluorescence (Fig. 5B, second panel; Movies S7; Fig 5C; Movie S8). In sharp contrast, neutrophils very efficiently engulfed and cleared large amounts of Δ*dotA*-GFP aggregated and trapped in the mesenchyme (Fig. 5B, lower panel, Movie S7) as well as low doses of WT-GFP (Fig 5B upper panel, Movie S7).

Altogether this shows that upon low dose bloodstream injection of *L. pneumophila*, macrophages and neutrophils efficiently cooperate to eliminate most of the injected bacteria within 20-24 hpi, with macrophages playing the primary role. However, *L. pneumophila* seems to persist upon high dose WT-GFP injection, and the observation of structures resembling large vacuoles containing live *L pneumophila* at 72hpi suggests that they can replicate in zebrafish macrophages. In contrast, neutrophils interact with *L. pneumophila* by quickly engulfing bacteria trapped in the mesenchyme near the site of injection but are less efficient in clearing blood-borne bacteria.

### Macrophages are the first line defence restricting L. pneumophila infection

In humans, innate immune responses, based essentially on the activity of professional phagocytes and the induction of pro-inflammatory cytokine genes, are the key players to control and restrict *L. pneumophila* proliferation. Hence, human disease develops primarily in immunocompromised individuals [10]. To investigate whether the phagocytes of the innate immune system, macrophages and neutrophils, are also responsible for controlling *L. pneumophila* infection in zebrafish larvae, we selectively and transiently depleted macrophages or neutrophils, and infected these “immunocompromised” larvae with *L. pneumophila*. Depletion of macrophages was achieved by knocking down the expression of *spi1b*, a transcription factor involved in early myeloid progenitor formation. A low dose of *spi1b* morpholino was reported to impact macrophages without affecting neutrophils [33]. We monitored the effect of low doses *spi1b* morpholino injection on macrophage and neutrophil populations in double transgenic larvae with green neutrophils (*mpx*:GFP) and red macrophages (*mfap4*:mCherryF). The specific depletion of macrophages was confirmed by counting macrophages and neutrophils at 72hpf (Fig S7A).

We then infected macrophage depleted larvae (*spi1b* knockdown) by intravenous injection of LD or HD of WT-GFP. Regardless of the infection dose, a dramatic decrease in larvae survival was observed, as even injection of low doses of WT-GFP resulted in the death of 30% of the larvae (Fig 6A). When injecting high doses of WT-GFP nearly all the infected larvae died by 72hpi, with the earliest deaths starting 48hpi (Fig 6A). In contrast, *spi1b* knockdown larvae injected with high doses of Δ*dotA*-GFP did not show impaired survival (Fig 6A). The increased mortality correlated with an increased but not significantly different bacterial burden in the *spi1b* knockdown larvae compared to control larvae (Fig 6B, Fig S2C). Intravital imaging of infected *spi1b* knock down larvae also showed that both low and high doses of WT-GFP failed to be cleared and that the bacteria established a replicative niche in the yolk, where they proliferated extensively (Fig 6C). This highlights, that macrophages are critical to restrict the onset of infection and *L. pneumophila* proliferation *in vivo*. Furthermore, these results also suggest that neutrophils, which are not depleted in *spi1b* knockdown larvae, fail to control *L. pneumophila* infection in the absence of macrophages.

**Figure 6.**
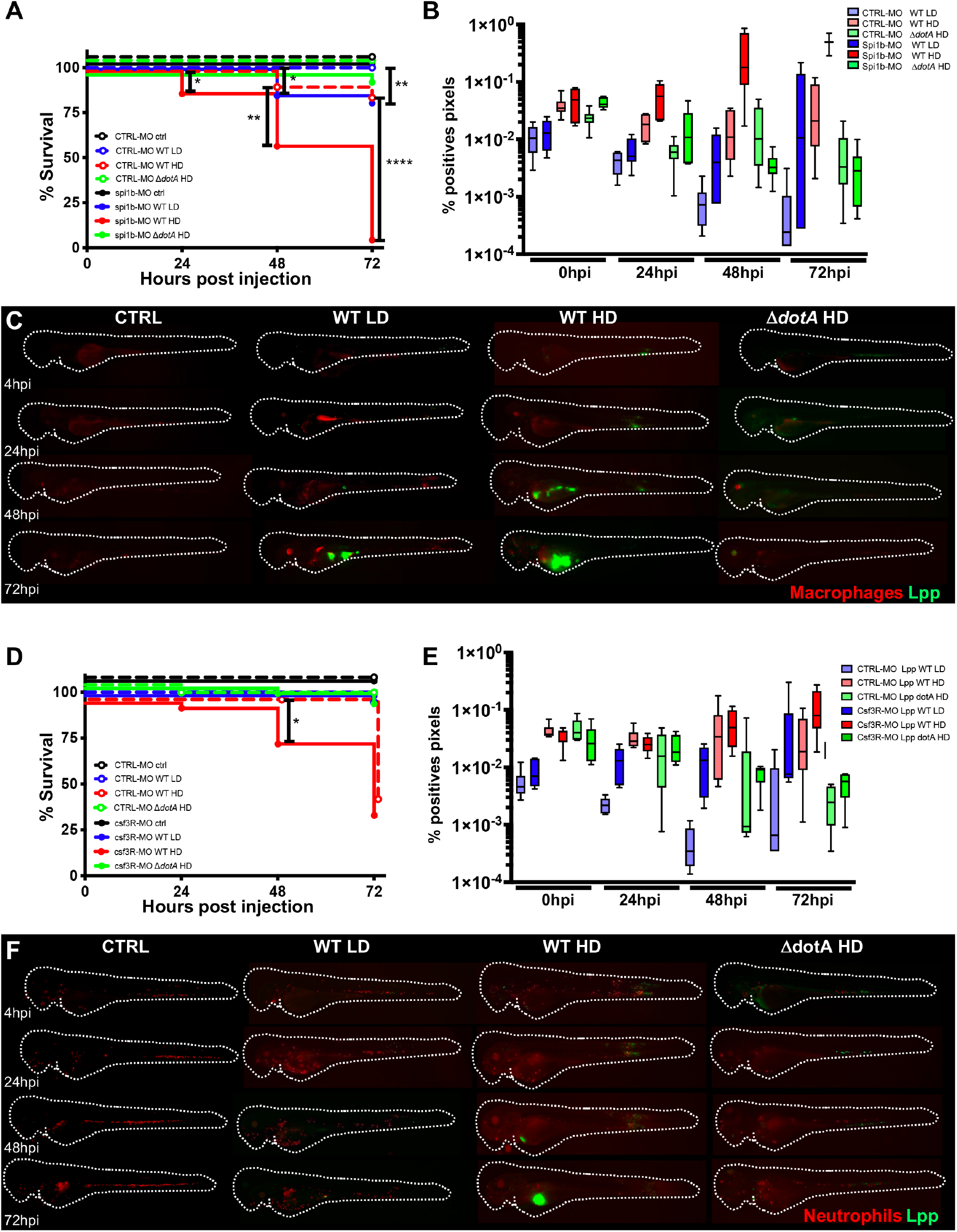
Macrophages are crucial to restrict *Legionella pneumophila* dissemination. **A)** Survival curves of CTRL morphant zebrafish larvae injected with a Low Dose (LD) (blue dashed curve, n=34 larvae) or a High Dose (HD) (red dashed curve, n=34) of WT-GFP, or with a HD (green dashed curve, n=24) of Δ*dotA* -GFP, and *spi1b* morphant zebrafish larvae injected with a LD (blue curve, n=48) or a HD (red curve, n=48) of WT-GFP, or with a High Dose (HD) (green curve, n=48) of Δ*dotA* -GFP. Non-injected CTRL morphant fish (black dashed curve, n=48), and *spi1b* morphant fish (black curves, n=48) were used as control. Infected and control larvae were incubated at 28°C. Data plotted are from two pooled independent experiments. **B)** and **E**) Bacterial burden evaluation by quantification of % of fluorescent pixel counts on individual injected larvae followed overtime by 0 to 72hpi. Each larva was imaged daily, and images were analysed with Fiji for bacterial burden evaluation. One experiment per phagocyte type plotted, 6 larvae for each condition. **D)** Survival curves of CTRL morphant zebrafish larvae injected with a LD (blue dashed curve, n=36) or a HD (red dashed curve, n=36) of WT-GFP, or with a HD (green dashed curve, n=24) of Δ*dotA* -GFP, and *csf3r* morphant zebrafish larvae injected with a LD (blue curve, n=24) or a HD (red curve, n=36) of WT-GFP, or with a HD (green curve, n=36) of Δ*dotA* -GFP. Non-injected CTRL morphant fish (black dashed curve, n=48), and *csf3r* morphant fish (black curve, n=36) were used as control. Data plotted are from two pooled independent experiments. Significant differences are highlighted with stars (see experimental procedure for statistical analysis) **C)** and **F)** Representative images of *L. pneumophila* dissemination, determined by live imaging using a fluorescence stereomicroscope, of Tg(*mfap4::mCherryF) spi1b* morphant larvae (**C**) and of Tg(*LysC::DsRed)^nz50^* (**F)** *csf3r* morphant larvae non infected, or infected with a LD or a HD of WT-GFP, or a HD of Δ*dotA*-GFP. The same infected larvae were live imaged 4h, 24h, 48h, and 72h post *L. pneumophila* injection. Overlay of GFP and mCherry fluorescence is shown. P < 0.05 was considered statistically significant (symbols: **** p < 0.0001; ***P < 0.001; **P < 0.01; *P < 0.05). No symbol on graphs means that not statistically differences were observed.

To analyse the role of neutrophils in controlling the infection, neutrophil development was disrupted by knocking down the G-CSF/GCSFR pathway using *csf3R* morpholino, previously reported to decrease up to 70% of the neutrophils present [34–36]. We monitored the efficiency of the *csf3R* morpholino knockdown in double transgenic larvae confirming that 75% of the neutrophil population was depleted, while macrophage numbers were only slightly decreased (Fig S7B). When HD Δ*dotA*-GFP was injected, neutrophil-depleted larvae survived, and the bacterial burden remained unchanged, like what we had observed in infections of macrophage-depleted larvae (Fig. 6D, E). However, when neutrophil-depleted larvae were injected with HD WT-GFP, larvae survival significantly decreased and bacterial burdens slightly increased by 48hpi (Fig. 6D, E; FigS2D). Intravital imaging showed that *csf3R* knockdown larvae that were unable to control *L. pneumophila* infection showed bacterial proliferation in the yolk comparable to WT control larvae (Fig 6F).

These results show that both macrophages and neutrophils are required for restricting and controlling *L. pneumophila* infection in the zebrafish model, but macrophages play the main role. Although neutrophils contributed less to clear the bacteria upon bloodstream injection, neutrophils might impact the infection outcome through cytokine release that can modulate macrophage activity.

### Key pro-inflammatory cytokine genes are induced upon L. pneumophila infection of zebrafish larvae

Proinflammatory cytokines produced by infected and bystander cells during *L. pneumophila* infection of humans and mice play crucial roles in orchestrating host defences to control infection [37, 38]. Infected cells produce IL-1α and IL-1β through a mechanism involving MyD88-dependent translational bypass. In contrast, bystander cells produce IL-6, TNF-a and IL-12 in an IL-1 receptor (IL-1R) dependant way [37, 39]. To determine the pro-inflammatory responses of zebrafish larvae during *L. pneumophila* infection, we analysed *il1b, tnfa*, and *ifng1/2* (orthologues of mammalian *Ifng*) gene expression levels over time by qRT-PCR on RNA isolated from individual infected larvae. We found that infection of zebrafish larvae with LD or HD of WT-GFP induced a rapid (by 6hpi) and robust induction of *il1b* and *tnfa* gene expression. In larvae injected with low doses of WT-GFP the expression levels started to decrease by 24hpi, and gradually became undetectable at 72hpi. In contrast, larvae injected with HD of WT-GFP, expression of *il1b* and *tnfa* did not decrease over time (Fig. 7A and B) and a significant induction of *ifng1* was observed at 48hpi (Fig. 7C) but not of *ifng2* (Fig. 7D). In parallel, we scored the bacterial burden of the infected larvae before the measurement of pro-inflammatory cytokine gene induction at each time point under the microscope, which consistently showed that larvae with increased *il1b* and *tnfa* induction had also high bacterial burdens in the yolk and were not controlling the infection (Fig S8A). These pro-inflammatory responses were T4SS dependent, as zebrafish larvae infected with HD of Δ*dotA*-GFP did not show significant induction of transcription of *tnfa, il1b* and *ing1/2* (Fig. 7 A-D).

**Figure 7.**
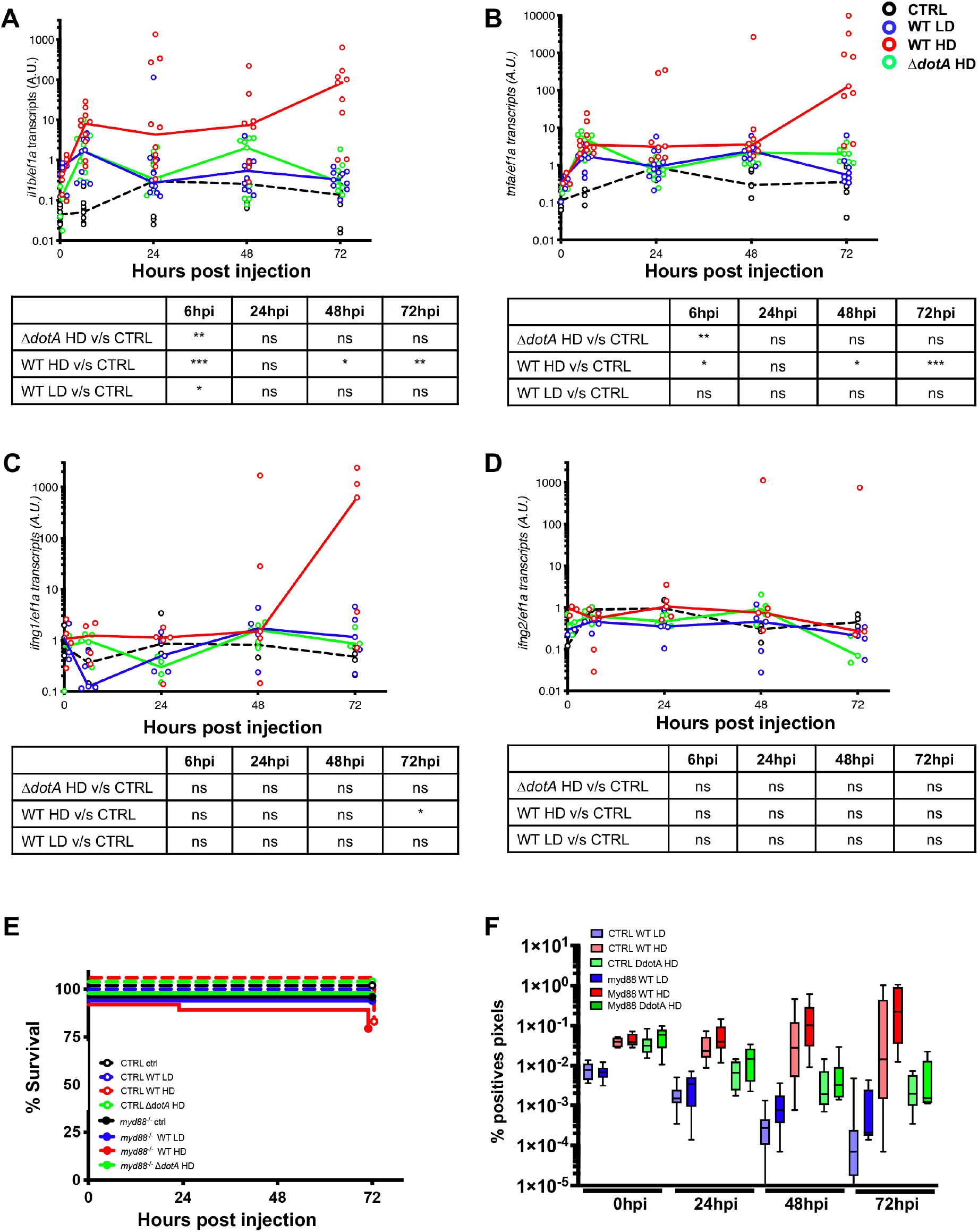
Cytokine gene induction upon *L. pneumophila* infection and Zebrafish larva Immunity to *L. pneumophila* is independent from signalling through MyD88 or compensated by other signalling pathways. **A-D)** Cytokine gene (*il1b, tnfa, ifng1, ifng2*) induction was measured from non-injected larvae as control (CTRL, dashed black curves) and individual zebrafish larvae injected with a LD (blue curves) or a HD (red curves) of WT-GFP, or a HD of Δ*dotA*-GFP (green curves). Data plotted are from 2 pooled experiments (n=10 larvae for each condition) for *il1b* and *tnfa*, and from 1 experiment (n=5 larvae for each condition) for *ifng1* and *ifng2;* individual values are shown, and curves correspond to the medians. Statistical analyses are shown in the table under each graph. **E)** Survival curves of CTRL zebrafish larvae injected with WT-GFP Low Dose (LD) (blue dashed curve) or High Dose (HD) (red dashed curve), or with Δ*dotA* -GFP HD (green dashed curve), and *myd88^hu3568^* mutant zebrafish larvae injected with WT-GFP LD (blue curve) or HD (red curve), or with Δ*dotA* -GFP HD (green curve). Non-injected CTRL larvae (black dashed curves), and *myd88^hu3568^* mutant larvae (black curves) were used as control. Infected and control larvae (n= 72 fish for *myd88^hu3568^* mutant conditions and n= 57 fish for CTRL conditions) were incubated at 28°C. Data plotted are from 3 pooled independent experiments. **F)** Bacterial burden evaluation by quantification of % of fluorescent pixel counts on individual injected larvae followed overtime from 0 to 72hpi. Each larva was imaged daily, and images were analysed with Fiji for bacterial burden evaluation. Two pooled experiments, 8 larvae for each condition. P < 0.05 was considered statistically significant (symbols: **** P < 0.0001; ***P < 0.001; **P < 0.01; *P < 0.05). No symbol on graphs means that not statistically differences were observed.

Collectively, these results reveal, that key pro-inflammatory cytokines known to orchestrate the host response during *L. pneumophila* infection in humans are also induced in zebrafish larvae, and that cytokine gene induction is sustained when uncontrolled *L. pneumophila* proliferation occurs.

### The immune response of zebrafish larvae to L. pneumophila infection is independent of MyD88 signalling

In innate immunity, the myeloid differentiation factor 88 (MyD88) plays a pivotal role in immune cell activation through Toll-like receptors (TLRs). MyD88-deficient mice are highly susceptible to *L. pneumophila* infection [40–43], however this is not the case when human macrophages are depleted of MyD88 [44]. Therefore, we sought to analyse which role MyD88 plays in zebrafish larvae during *L. pneumophila* infection. We injected *myd88-/-* and control larvae with LD or HD of WT-GFP, or with HD of Δ*dotA-*GFP and monitored larvae survival and bacterial burden over time as described in Figure 1. Our results show that susceptibility to infection of *myd88*-/- larvae injected with HD of WT-GFP was comparable to that of WT larvae, as only a very slight, but no significant increase of the bacterial burden was observed in *myd88*-/- larvae by 24hpi. *L. pneumophila ΔdotA-GFP* did not establish an infection, and the bacterial burden decreased over time indicating that bacteria were cleared (Fig. 7E, F; Fig S2E). To determine if pro-inflammatory responses were affected in the absence of MyD88 signalling, we analysed *il1b, tnfa* and *ing1/2* gene expression levels over time in control and *myd88*-/- larvae. Our results showed that *il1b, tnfa* and *ing1/2* gene expression levels were comparable in control and *myd88*-/- infected larvae for all tested conditions (LD WT-GFP and HD Δ*dotA*-GFP (Fig S8B, C).

Taken together, our results suggest that MyD88 signalling is not required for the innate immune response against *L. pneumophila* infection in the zebrafish larvae, similar to what is seen in human cell infection models. However, MyD88 signalling may also be functionally compensated by other immune signalling pathways.

## DISCUSSION

In this study, we developed a zebrafish larva infection model for *L. pneumophila* and have analysed host pathogen interactions as well as the host innate immune response to infection. We have found that a successful infection of zebrafish larvae by *L. pneumophila* depends on the infection site, the infection dose, the T4SS Dot/Icm and the innate immune response of the host, in particular on the action of macrophages and neutrophils. Wild type zebrafish larvae are susceptible to infection in a dose dependent manner, as larvae injected in the bloodstream with increasing doses of bacteria developed an infection with bacterial dissemination and replication, concomitant with host death proportional to the initial injected bacterial load. However, the fact that only about 30% of the larvae displayed this phenotype, indicates that the innate immune defence of the larvae against *L. pneumophila* infection is relatively efficient. Thus, the establishment of infection is determined not only by the infection dose but also by the capacity of the host immune system to quickly and efficiently fence off the infection.

Like in several other zebrafish infection models, only blood borne bacteria can proliferate and induce mortality in zebrafish larvae (Emma do you have some papers we could cite?). Once in the blood circulation, bacteria are engulfed and are eliminated by both macrophages and neutrophils in a dose-dependent manner. However, T4SS competent *L. pneumophila* are also able to reach the yolk region, cross the endothelium of the yolk vessels and enter the yolk sac region. Once there, *L. pneumophila* gains a significant advantage in the pathogen-host arms race and establishes a replicative niche where it proliferates extensively, forming complex, aggregated bacterial structures, located below and above the yolk cell membrane, that professional phagocytes fail to clear. In the yolk sac region *L. pneumophila* is protected from the host immune system, when it reaches the yolk cell, as professional phagocytes are unable to enter in the yolk. Extensive proliferation of the bacteria eventually leads to host death, likely due to exhaustion of the nutrients present in the yolk, which are key in supporting the larvae development. Interestingly, we have also observed that in few cases (less than 5%) the infected larvae were able to extrude the bacterial aggregates growing in the yolk and survived. This host defence mechanism has also been reported in a caudal fin model of *Mycobacterium marinum* infection, where infected zebrafish larvae extruded the bacteria-containing granuloma and during infection with *Aspergillus fumigatus* [45, 46].

To our knowledge, the tropism, and the establishment of a replicative niche in the yolk upon pathogen injection in the bloodstream is a unique feature of *L. pneumophila*. Our results have shown that, when macrophages and neutrophils are depleted, blood-borne *L. pneumophila* can still reach the yolk cell. It seems that *L. pneumophila* can cross the venous endothelium of the yolk and reach the nutrient-rich content of the yolk cell. Interestingly, direct yolk cell injection revealed that also only the WT but not the T4SS knockout strain is able to replicate and establish a persistent infection in the yolk, irrespective of the dose injected. This result points towards the involvement of the T4SS system and its secreted effectors in infection and replication but also nutrient uptake in the yolk environment. Yolk sac injection has already been used previously for the analyses of *Candida albicans* infection or the study of streptococcal infections, however in this case the pathogens disseminated from the yolk sac into the animal [47, 48], a phenotype not observed when *L. pneumophila* was injected in the yolk sac. Further analyses of the phenotype of yolk sac replication in embryonated chicken eggs showed again that only WT *L. pneumophila* can replicate in the yolk region, further supporting the importance of the T4SS in nutrient uptake in addition to its known role in infection (Fig. S5A, B, C). *L. pneumophila* is known to mainly use amino acids as carbon and energy sources for growth [49] and secreted T4SS effectors have been shown to aid in amino acid uptake [50]. However, fatty acids, glucose and/or glycerol also serve as carbon sources during the later stages of the life cycle of *L. pneumophila* [51, 52], but effectors connected to the uptake of these nutrients have not been identified yet. The yolk cell is composed of a complex and dynamic mixture of different lipids on which the zebrafish larvae rely on for nutrition throughout development in the early larva phase. Cholesterol and phosphatidylcholine are the main constituents until 120hpf, with triacylglycerol, phosphatidylinositol, phosphatidylethanolamine, diacyl-glycerol, cholesteryl esters and sphingomyelins also present in significant concentrations [53]. *L. pneumophila* is known to secrete several effectors with lipolytic activity through its T4SS which could be important for growth in a lipid rich environment like the yolk [54]. In a first attempt to identify one of these effectors we analysed the growth of a *L. pneumophila* mutant for a gene encoding a sphingosine-1 phosphate lyase (*Lp*Spl) [27]. When compared to the WT strain after direct injection in the zebrafish larvae yolk sac a small but significant difference in larvae mortality was observed for the Δ*spl* strain, suggesting that *Lp*Spl is one of several effectors that might participate in nutrient acquisition from lipids (Fig. S5A). However, further analyses are needed to characterize the involvement of *Lp*Spl in nutrient acquisition during infection of the yolk region and to identify other effectors potentially implicated in this phenotype.

Studies of *Legionella* infection in humans, guinea pigs and mouse lungs have shown that *L. pneumophila* interacts closely with neutrophils and mononuclear phagocytes [55, 56]. Professional phagocytes are the main replication niche for *L. pneumophila* with monocytes and macrophages, and in particular alveolar macrophages, representing the main cells for replication in the lungs [57–60]. *In vivo* studies in mice have shown that upon lung infection with *L. pneumophila* neutrophils, cDCs, monocytes, and monocyte-like cells are rapidly recruited to the infection site, but although all these cells seem to engulf the bacteria, *L. pneumophila* can only translocate effectors into neutrophils and alveolar macrophages. In zebrafish macrophages appear during the first days of development, followed by neutrophils a day later, with both forming an efficient immune system that protects the developing fish [25, 61–63]. Therefore, the zebrafish larva offers a unique possibility to interrogate the role of innate immune responses to infection [23]. Indeed, macrophage depleted larvae showed a dramatically increased susceptibility to *L. pneumophila* infection as nearly 100% of larvae inoculated with HD of WT and 30% of larvae inoculated with LD of *L. pneumophila* died from the infection. Hence, macrophages are the first line of infection control against *L. pneumophila* and are essential for restricting and controlling blood-borne infections, similar to what was observed for *Burkholderia cenocepacia or Staphylococcus aureus* zebrafish infection [64, 65]. In contrast, when neutrophils were depleted, the innate immune response was less affected, suggesting that neutrophils are also required to ensure an effective innate immune response and, that macrophages alone are not able to contain high burdens of *L. pneumophila* infection (Fig. 6).

Human innate immune signalling relies strongly on activation of Toll-like receptors (TLRs) and respective adaptor molecules, all of which are highly conserved in the zebrafish [66, 67]. One of these adaptors is MyD88, known as a central player in interleukin 1 receptor (IL-1R) and TLR signalling in humans and mammalian models [68]. MyD88 signalling is crucial for mice to combat *L. pneumophila* infection, as it triggers the early secretion of inflammatory cytokines, neutrophil recruitment, and the host immune response to the infection. Consequently, mice that lack MyD88 are highly susceptible to infection [39–42]. However, in MyD88 depleted human macrophages *L. pneumophila* replication is not different to replication in WT cells [44] Here we show, Myd88 signalling does not play a crucial role or may be redundant in the control of the innate immune response to *L. pneumophila* infection of zebrafish larvae, suggesting that zebrafish infection mirrors human infection better than the mouse model. In the mouse model infected macrophages are incapable of producing cytokines, such as tumour necrosis factor (TNF) and interleukin-12 (IL-12), which are necessary to control infection. In contrast, infection of zebrafish larvae with WT *L. pneumophila* induced a rapid (by 6hpi) and robust induction of *il1b* and *tnfa* gene expression. However, it is thought that IL-1 released initially by *L. pneumophila-infected* macrophages drives the production of critical cytokines by bystander cells [37]. Infection of zebrafish larvae with HD of WT *L. pneumophila* induced a rapid (by 6hpi) and robust induction of *il1b* and *tnfa* gene expression whereas WT LD infection leads only to a short induction of *Il1b* transcript levels at 6hpi before declining to CTRL levels at later time points, suggesting that a short boost of IL-1β is sufficient to control LD of *L. pneumophila*. However, for a high load of *L. pneumophila* even a high and long-term induction of IL-1β is not sufficient to warrant infection control, suggesting that the self-regulation of the immune response may be abrogated leading to a constant activation of IL-1β expression. Moreover, gene expression analyses also confirms that Myd88 has no influence on the control of the inflammatory response, as no statistically significant difference in the transcript levels of *il1b, tnfa, ifng1 or infg2* was observed further suggesting that activation of the IL1R and certain TLR pathways are not crucial for *L. pneumophila* clearance in zebrafish larvae. It is even possible that IL-1β release is beneficial for *L. pneumophila* replication, as it was shown that IL-1β may also indicate an activation of the metabolic state of the bystander cells as it was shown that IL-1β induces a shift towards more metabolically active cells and increased cellular glucose uptake [69], which could aid *L. pneumophila* replication.

In conclusion, the zebrafish infection model for *L. pneumophila* mimics the immune response known of human infection and recalls the essentiality of the type IV secretion system for virulence of this pathogen. The unique advantages of the zebrafish model provide now exciting possibilities to further explore *L. pneumophila* host interactions and to interrogate at the molecular level the bacterial determinants and host factors involved in the dynamics of bacterial dissemination, the molecular basis of yolk region invasion and the interactions of the bacteria with macrophages and neutrophils.

## MATERIALS AND METHODS

### Ethics Statement

Animal experiments were performed according to European Union guidelines for handling of laboratory animals (http://ec.europa.eu/environment/chemicals/lab_animals/home_en.htm) and were approved by the Institut Pasteur Animal Care and Use Committee. and the French Ministry of Research (APAFIS#31827). The inoculation of embryonated chicken eggs is a standard procedure in diagnostics for multiplication and antigen production of *Legionella* and is not covered by the national law for animal experiments in France (Décret n° 2013-118 du 1er février 2013).

### Zebrafish care and maintenance

Wild-type AB fish, initially obtained from the Zebrafish International Resource Center (Eugene, OR), Tg(*Lyz::DsRed)*^nz50^ [70], Tg*(mfap4::mCherryF*) (ump6Tg) [36] Tg(mpx:GFP)^i114^ [71], Tg(kdrl::mCherry)^is5^ [72] and *myd88^hu3568^* mutant line (obtained from the Hubrecht Laboratory and the Sanger Institute Zebrafish Mutation Resource) [73], were raised in our facility. Eggs were obtained by natural spawning, bleached according to standard protocols, and kept in Petri dishes containing Volvic source water and, from 24 hours post fertilization (hpf) onwards 0.003% 1-phenyl-2-thiourea (PTU) (Sigma-Aldrich) was added to prevent pigmentation. Embryos were reared at 28°C or 24°C according to the desired speed of development; infected larvae were kept at 28°C. Timings in the text refer to the developmental stage at the reference temperature of 28.5°C. Larvae were anesthetized with 200μg/ml tricaine (Sigma-Aldrich) during the injection procedure as well as during *in vivo* imaging and processing for bacterial burden evaluation or cytokine expression studies.

### Bacterial strains and growth conditions

*Legionella pneumophila* strain Paris carrying the pNT28 plasmid encoding for green fluorescent protein (constitutive GFP) [74], wild-type (WT-GFP) or Δ*dotA*-GFP were plated from −80°C glycerol stocks on N-(2-acetamido)-2-aminoethanesulfonic acid (ACES)-buffered charcoal yeast-extract (BCYE) medium supplemented with 10 μg/ml of chloramphenicol and cultured for 3 days at 37°C. Suspensions were prepared by resuspending bacteria in sterile 1x Phosphate Buffered Saline (PBS) and adjusting the OD 600 according to the desired bacterial concentrations for injection.

### Morpholino injections

Morpholino antisense oligonucleotides (Gene Tools LLC, Philomath, OR, USA) were injected at the one to two cell stage as described [75]. A low dose (4ng) of *spi1b* (previously named pu1) translation blocking morpholino (GATATACTGATACTCCATTGGTGGT) [76] blocks macrophage development only but can also block neutrophil development when it is injected at a higher dose (20ng in 2nl). The csf3r translation blocking morpholino (GAACTGGCGGATCTGTAAAGACAAA) (4ng) [34] was injected to block neutrophil development. Control morphants were injected with 4ng control morpholino, with no known target (GAAAGCATGGCATCTGGATCATCGA).

### Amoebae infections

*Acanthamoebae castellanii* was infected with *L. pneumophila* wild type expressing GFP and then used to assess if zebrafish develop an infection when ingesting infected amoebae. *A. castellanii* was seeded in a flask with infection buffer (4 mM MgSO_4_, 0.4 M CaCl_2_, 0.1% sodium citrate dihydrate, 0.05 mM Fe(NH_4_)2(SO_4_)2×6H_2_O, 2.5 mM NaH_2_PO_3_, 2.5 mM K_2_HPO_3_), and after 1 hour of attachment the cells were infected with the bacteria at MOI 1. After one hour of infection, the amoebae were washed three times with PBS and fresh infection buffer was added. After 48 hours of infection, the amoebae were carefully washed, detached, and used for the bath immersion experiment.

### Bath immersion

Experiments were performed using the AB or Tg(mfap4::mCherryF) zebrafish lines maintained at 28 °C under standard conditions in Volvic water. Bath immersion infections were done with 120hpf larvae that already had a developed swim bladder. Groups of 10 larvae were distributed into 6-well plates containing 4.0 ml/well of Volvic spring water and either 1ml of bacterial suspension, 1ml of *L. pneumophila-containing* amoebae, or 1ml of non-infected amoebae, all in PBS. The plates were incubated at 28 °C for 24 hours, and then larvae were individually distributed in an individual well in 24 wells culture plates and monitored by imaging using a fluorescence stereomicroscope.

### Zebrafish injection

The volume of injected suspension was deduced from the diameter of the drop obtained after mock microinjection, as described in [75]. Bacteria were recovered by centrifugation, washed, resuspended at the desired concentration in PBS. 72h post-fertilization (hpf) anesthetized zebrafish larvae were microinjected iv or in closed cavities, or the yolk with 0.5-1nl of bacterial suspension at the desired dose (~10^3^ bacteria/nl for Low Dose (LD) and ~10^4^ bacteria/nl for High Dose (HD) as described [24, 32]. Infected larvae were transferred into individual wells (containing 1ml of Volvic water + 0.003% PTU in 24-well culture plates), incubated at 28°C and regularly observed under a stereomicroscope, twice a day overtime up to 72hpi.

### Evaluation of the bacterial burden in infected larvae

The bacterial burden was measured routinely by counting the total number of fluorescent pixels corresponding to the GFP channel using the Metavue software 7.5.6.0. Briefly, images corresponding to the GFP channel were adjusted to a fixed threshold that allowed to abrogate the background of the autofluorescence of the yolk. The same threshold was used for all images. The histogram in the “Analyze” menu was used to obtain the number of black and white pixels. As shown in figure S1A, the percentage of white pixels in each image corresponding to *L. pneumophila* was plotted using GraphPad Prism^®^ software. This method was chosen to routinely quantify bacterial burden as it allows to follow each infected larva overtime (individual larvae were imaged with the fluorescence stereomicroscope daily using the same settings from 0 to 72hpi).

### Bacterial burden analyses by FACS

Injected zebrafish larvae were collected at 0, 24, 48 and 72hpi and lysed. Each larva was placed in a 1.5 ml Eppendorf tube and anesthetized with tricaine (200μg/ml), washed with 1ml of sterile water and placed in 150 μl of sterile water. Larvae were then homogenized using a pestle motor mixer (Argos). Each sample was transferred to an individual well of a 96 well plate, counted on a MACSQuant VYB FACS (Miltenyi Biotec) and data analysed using FlowJo version 7.6.5. as shown in Figure S1B. For CFU enumeration, serial dilutions were plated on BCYE agar plates supplemented with10ug/ml chloramphenicol and the *Legionella* Selective Supplement GVPN (Sigma) added according to the manufacturer’s instructions. Plates were incubated for 4-5 days at 37°C and colonies with the appropriate morphology and colour were scored using the G-Box imaging system (Syngene) and enumerated using the Gene Tools software (Syngene) as shown in Figure S1C. Manual quantification was also performed to identify absolute absence or presence of bacteria in the different zebrafish compartments. Larvae with a single GFP dot (indicating the presence of few bacteria) were considered as infected. The resulting statistical presence map was used to follow the evolution of the infection and dissemination of *L. pneumophila* in zebrafish larvae over time (Figure 1E).

### Inoculation and quantification of *L. pneumophila* strains in *in ovo* experiments

Fertilized chicken eggs purchased from a local producer (Saint-Maurice-sur-Dargoire, Rhône, France) were incubated at 35°C in an egg incubator (Maino, Italy) to maintain normal embryonic development. Eggs were pathogen and antibiotic free. On day 0, 23 embryonated chicken eggs (ECE) were inoculated at 8 days of embryonation (DOE) with either *L. pneumophila* WT (n=9), *L. pneumophila ΔdotA* (n=7) or sterile PBS as control (n=7). *L. pneumophila* concentration in WT and Δ*dotA* suspensions before ECE injection was quantified at 9.2 log_10_ CFU/mL and 9.1 log_10_ CFU/mL, respectively. *L. pneumophila* concentration in the yolk sac of ECE directly after injection were estimated, considering both the measured inoculum counts and the yolk sac volumes (median (interquartile range) [IQR] volume, 30 [28.7-31.2] mL), at 7.4 and 7.3 log_10_ CFU/mL in the WT and Δ*dotA* groups, respectively. Two-day cultures of Lpp-WT and Lpp-Δ*dotA* on BCYE at 36°C were suspended in PBS at a DO = 2.5 McFarland (9 log_10_ CFU/mL) and 0.5 mL of suspensions or PBS (negative control) were inoculated in the yolk sac of ECE. After inoculation, ECE were candled every 24 hours to assess embryo viability until day-6 post infection. Embryos that died the day after inoculation (n=2, corresponding to one WT-infected and one Δ*dotA*-infected embryo) were discarded for *L. pneumophila* quantification as death was probably due to bad inoculation. Dead embryos were stored at 4°C overnight prior to harvesting the yolk sacs. Remaining live embryos at 6-days post injection were euthanized by overnight refrigeration and the yolk sacs were collected. After measuring their volume, yolk sacs were homogenized using gentleMACS™ Octo Dissociator (Miltenyi Biotec, Germany) and 100 μL of serial dilutions at 10^-2^, 10^-4^ and 10^-6^ were automatically plated using easySpiral^®^ automatic plater (Interscience, France) in triplicates on BCYE agar. *L. pneumophila* were quantified after 5 days-incubation using Scan^®^ 1200 Automatic HD colony counter (Interscience, France). Comparison of survival curves was performed using Log-rank (Mantel-Cox) test. P < 0.05 was considered statistically significant. Comparison of *L. pneumophila* quantifications between WT- and Δ*dotA*-infected embryos was done using Mann-Whitney test. P < 0.05 was considered statistically significant (symbols: **** P < 0.0001; ***P < 0.001; **P < 0.01; *P < 0.05).

### Live imaging, image processing and analysis

Quantification of total neutrophils and/or macrophages on living transgenic reporter larvae upon infection was performed as previously described [32]. Briefly, bright field, DsRed and GFP images of whole living anesthetized larvae were taken using a Leica Macrofluo™ Z16 APOA (zoom 16:1) equipped with a Leica PlanApo 2.0X lens, and a Photometrics^®^ CoolSNAP™ *HQ2* camera. Images were captured using Metavue software 7.5.6.0 (MDS Analytical Technologies). Then larvae were washed and transferred in a new 24 wells plate filled with 1ml of fresh Volvic water per well, incubated at 28°C and imaged again under the same conditions the day after. Pictures were analysed, and Tg(*lyzC::DsRed*) neutrophils or Tg(*mfap4::mCherryF*) macrophages manually counted using the ImageJ software (V 1.52a). Counts shown in figures are numbers of cells per image. The quantification of fluorescence images was also done using CellProfiler™ Software (Broad Institute) using two semi-automatic batch pipelines. Both pipelines normalize the intensity, operate image pre-processing, and use thresholding to calculate the percentage of area positive for macrophage/neutrophils and bacteria, normalized on the whole image area. Both pipelines also take advantage of manual editing to increase identification accuracy and define the yolk area. The positive signal is then automatically masked to calculate the amount of signal in the body or yolk of each zebrafish for all the experiments and produce a .csv file used for subsequent statistical treatment (Figure 3F and Figure S6). High resolution confocal live imaging of injected larvae was performed as previously described [77]. Briefly, injected larvae were positioned in lateral or ventral position in 35 mm glass-bottom-Dishes (Ibidi Cat#: 81158) or in glass bottom-8well-slides (Ibidi Cat#: 80827). Larvae were immobilized using a 1% low-melting-point agarose (Promega; Cat#: V2111) solution and covered with Volvic water containing tricaine. A Leica SP8 confocal microscope equipped with two PMT and Hybrid detector, a 10X dry (PL Fluotar 10X dry:0.30), 20X IMM (HC PL APO CS2 20X/0.75), or a 40x water IMM (HC PL APO CS2 40X/1.10) objective, a X–Y motorized stage and with LAS-X software, was used to live image injected larvae. To generate images of the whole larvae, a mosaic of confocal z-stack of images was taken with the 10X or 20X objective using the Tile Scan tool of the LAS-X software and was stitched together using the Mosaic Merge tool of the LAS-X software. All samples were acquired using the same settings, allowing comparisons of independent experiments. The acquisitions were post processed with the Lightning tool of the LAS-X software to eliminate noise (deconvolution). After acquisition, larvae were washed and transferred in a new 24-well plate filled with 1 ml of fresh Volvic water per well, incubated at 28°C and imaged again under the same conditions over time. A Leica SPE inverted confocal microscope and a 40x oil IMM objective (ACS APO 40 × 1.15 UV) was also used to live image larvae infected with *L. pneumophila ΔdotA-GFP* (Fig 4, 5). The 3D or 4D files generated by the time-lapse acquisitions were processed, cropped, analysed, and annotated using the LAS-X and LAS-AF Leica software. Acquired Z-stacks were projected using maximum intensity projection and exported as AVI files. Frames were captured from the AVI files and handled with Miocrosoft PowerPoint ^®^ (Microsoft Office 365) software to mount figures. AVI files were cropped and annotated using ImageJ software. Files generated with the LAS-X software were also processed and analysed with the Imaris software version9.5 (Bitplane, OXFORD Instruments) for 3D or 4D reconstruction, surfacing and volume rendering. The yolk region has been manually annotated (Movie S5).

### qRT-PCR to measure gene expression of cytokine encoding genes

RNA was extracted from individual larvae using the RNeasy^®^ Mini Kit (Qiagen). cDNA was obtained using M-MLV H-reverse-transcriptase (Promega) with a dT17 primer. Quantitative PCR was performed on an ABI7300 thermocycler (Applied Biosystems) using TakyonTM ROX SYBR^®^ 2X MasterMix (Eurogentec) in a final volume of 10 μl. Primers used: *ef1a (housekeeping gene used for normalization):* GCTGATCGTTGGAGTCAACA and ACAGACTTGACCTCAGTGGT*; il1b:* GAGACAGACGGTGCTGTTTA and GTAAGACGGCACTGAATCCA*; tnfa:* TTCACGCTCCATAAGACCCA and CAGAGTTGTATCCACCTGTTA*; ifng-1-1:* ACCAGCTGAATTCTAAGCCAA and TTTTCGCCTTGACTGAGTGAA; *ifng-2:* GAATCTTGAGGAAAGTG AGCA and TCGTTTTCCTTGATCGCCCA

### Statistical analysis

Normal distributions were analysed with the Kolmogorov–Smirnov and the Shapiro–Wilk tests. To evaluate difference between means of normally distributed data (for neutrophil and macrophage numbers), an analysis of variance followed by Bonferroni’s multiple comparison tests was used. For bacterial burdens (CFU/FACS counts), values were Log10 transformed. For cytokine expression and bacterial burdens (evaluated by fluorescent pixel count, or FACS) non-Gaussian data were analysed with the Kruskal–Wallis test followed by Dunn’s multiple comparison test. P < 0.05 was considered statistically significant (symbols: **** P < 0.0001; ***P < 0.001; **P < 0.01; *P < 0.05). No symbol on graphs means that not statistically differences were observed. Survival data were plotted using the Kaplan–Meier estimator and log-rank (Mantel–Cox) tests were performed to assess differences between groups. Statistical analyses were performed using GraphPad Prism^®^ software. Statistical analyses for *in ovo* experiments, were performed using GraphPrism version 9.4.

## Author contributions

FV, LB, SJ, ECG and CB designed the experiments. FV, LB, VL, DS, MI and ECG performed the experiments. FV, LB, VL, ECG analysed the experiments. FV drafted, and ECG and CB wrote the manuscript with input from all authors. ECG and CB supervised the work and acquired funds.

## Data Availability

All relevant data are within the paper and its Supporting Information files. The movies related to the paper are available at https://zenodo.org/deposit/7410466

## Competing Interest

The authors declare there are no competing interests.

## ACKNOWLEDGEMENTS

We thank Pedro Escoll and Jean-Pierre Levraud for help and ideas in the initial set up of the project and Tobias Sahr for critical reading and helpful discussions, Philippe Herbomel for helpful scientific advice, and Yohann Rolin for his excellent care of the fish. Work in the CB laboratory is financed by the Institut Pasteur, the Fondation pour la Recherche Médicale (FRM) grant EQU201903007847 and the grant ANR-10-LABX-62-IBEID and ANR-22-CE15 0009 01 ZebraLegion. Work in ECG group is financed by Institut Pasteur, CNRS, and ANR 17-CE15-0026 and 20-CE15-0024 and ANR-22-CE15 0009 01 ZebraLegion. Valerio Laghi is funded by ANR 20-CE15-0024. We acknowledge the Image analyses Hub at Institut Pasteur for the use of the IMARIS workstation. The funders, other than the authors, did not play any role in the study or in the preparation of the article or decision to publish.

## Notes

https://zenodo.org/deposit/7410466

### Competing Interest Statement

The authors have declared no competing interest.

### Summary of Updates

Additional experiments and analyses have been udnertaken: - Bath immersion experiment where zebrafish are infected via a natural route such as Acanthamoeba castellani infected with Legionella pneumophila - Injection of L. pneumophila in additional closed cavities such as the otic vesicle and the hind brain - Live imaging at high resolution of Legionella-leukocytes interactions in the yolk region, mounted and added two new movies for better analyses of the infection phenotype. All high-resolution movies can be accessed at https://zenodo.org/record/7307933 - Re-analyses of all experiments to additionally quantify bacterial burdens based on the fluorescence pixel count of the bacteria in the larvae during infection, and to quantify % of infected larvae showing bacterial burden in the yolk region - Quantification of leukocyte recruitment to bacteria growing in the yolk - Better description of statistics and adding of statistics where it was missing

