## Supplemental data for "Hiding in the yolk: a unique feature of *Legionella pneumophila* infection of zebrafish"

**Figure S1:** Comparison of three methods to estimate the bacterial burden of infected zebrafish larvae

**Figure S2: Bacterial burden evaluation by FACS overtime on individual lysed larva after *Legionella pneumophila* infection.**

**Figure S3:** *Legionella pneumophila* invades the yolk only upon bloodstream inoculation and only blood borne *L. pneumophila* WT proliferate in the yolk region of zebrafish**.**

**Figure S4:** Bath immersion of 120 hpf zebrafish larvae using *Acanthamoeba castellanii* infected with wild type *L. pneumophila*

**Figure S5:** Survival curves of zebrafish larvae directly injected with a *L. pneumophila* WT, *Δspl* or a *ΔdotA* mutant in the yolk of zebrafish or chicken eggs

**Figure S6:** Quantification of the bacterial burden in the whole body, in the body or in the yolk region compared to the quantification of macrophages or neutrophils in the body or in the yolk region in HD WT-GFP infected larvae overtime

**Figure S7:** Macrophage and neutrophil depletion by morpholino: evaluation of the impact on the non-depleted leukocyte population

**Figure S8.** Correlation between bacterial burden (evaluated by fluorescence on individual injected larvae before RNA extraction) and cytokine gene induction at 48 and 72hpi upon bloodstream injection LD, HD WT or HD Δ*dot*A *L pneumophila* strain

**Supplementary Movies:**

**All supplementary movies are available at**
**https://zenodo.org/record/5572884#.YX0fHR1S-jR**

**Movie S1:** *L. pneumophila* growing in the yolk region at 72hpi: localization in the yolk in AB wild type larva (related to Figure 2D)

**Movie S2:** *L. pneumophila* growing in the yolk region at 72hpi: interactions with blood vessels (related to Figure 2E)

**Movie S3:** *L. pneumophila* growing in the yolk region at 72hpi: interactions with macrophages (related to Figure 3B)

**Movie S4:** *L. pneumophila* growing in the yolk region at 72hpi: interactions with neutrophils (related to Figure 3C)

**Movie S5:** *L. pneumophila* growing in the yolk region between 48 and 72hpi (related to Fig 3E)

**Movie S6:** Macrophage - *L. pneumophila* interactions (LD, HD WT and HD ⍰dotA) **(**related to Figure 4)

**Movie S7:** Neutrophil - *L. pneumophila* (LD, HD WT and HD ⍰dotA) interactions (related to Figure 5)

**Movie S8:** Dying phagocytosing neutrophils upon *L. pneumophila* HD injection (related to Figure 5)

**Figure legends**

**Figure S1: Comparison of three methods to estimate the bacterial burden of infected zebrafish larvae at different points post infection**. A) For bacterial burden measure by fluorescent pixel counts, the pictures corresponding to the GFP channel were analysed to quantify the percentage of fluorescent pixels using the ImageJ software. Individual larvae injected with WT-GFP Low Dose (LD) (blue symbols) or High Dose (HD) (red symbols) or infected with Δ*dotA*-GFP HD (green symbols) have been plotted and represented as box plot. Two independent experiments pooled, n=10 larvae per condition). **B**) For FACS analyses, infected larvae were lysed and then GFP bacteria were counted on a MACSQuant VYB FACS (Miltenyi Biotec). One experiment plotted, n=5 larvae per condition. **C**) CFUs were enumerated by plating serial dilutions of lysed infected larvae in BCYE agar supplemented with Chloramphenicol and *Legionella* Selective Supplement GVPN (Sigma). One experiment plotted, n=5 larvae per condition. P < 0.05 was considered statistically significant (symbols: **** P < 0.0001; ***P < 0.001; **P < 0.01; *P < 0.05). No symbol on graphs means that not statistically differences were observed.

**Figure S2: Bacterial burden evaluation by FACS overtime on individual lysed larva after *Legionella pneumophila* infection.** For FACS analyses, individual infected larvae were lysed and then GFP bacteria were counted on a MACSQuant VYB FACS (Miltenyi Biotec). **A)** Related to Fig 1C: 6 pooled experiments, n= 26 larvae for WT HD (28 for 72h), n= 26 larvae for WT LD (30 for 72h), n= 25 larvae for *ΔdotA* HD (26 for 72h) **B)** Related to Fig 2B: Fluorescent pixel count evaluation overtime upon yolk cell injection. One experiment is plotted. N= 6 larvae for WT HD, n= 6 larvae for WT LD, n= 5 larvae for *ΔdotA* HD**. C)** Related to Fig 6B: 2 pooled experiments, n= 8 Spi1b-MO larvae for WT HD, n= 8 Spi1b-MO larvae for WT LD, n= 5 Spi1b-MO larvae for *ΔdotA* HD, n=5 control larvae for WT HD, n=5 control larvae for WT LD and n=5 control larvae for *ΔdotA* HD. **D)** Related to Fig 6E: 2 pooled experiments, n= 8 Csf3R-MO larvae for WT HD (10 for 72h), n= 8 Csf3R-MO larvae for WT LD (9 for 72h), n= 5 Csf3R-MO larvae for *ΔdotA* HD (4 for 0h), n=6 control larvae for WT HD (8 for 72h), n=6 control larvae for WT LD (9 for 72h) and n=5 control larvae for *ΔdotA* HD (6 for 72h). **E)** Related to Fig 7B: 3 pooled experiments, n= 13 *myd88* larvae for WT HD, n= 13 *myd88* larvae for WT LD (12 for 0h), n= 13 *myd88* larvae for *ΔdotA* HD, n=10 control larvae for WT HD, n=10 control larvae for WT LD, n=10 control larvae for *ΔdotA* HD. P < 0.05 was considered statistically significant (symbols: **** P < 0.0001; ***P < 0.001; **P < 0.01; *P < 0.05). No symbol on graphs means that not statistically differences were observed.

**Figure S3: *L. pneumophila* invades the yolk only upon bloodstream inoculation and only blood borne *L. pneumophila* WT proliferate in the yolk region of zebrafish. A)** Scheme of 72hpf larva indicating the sites of bacterial injection. Site of injection are indicated by green dashed boxes. OV: otic vesicle; HBV: hind brain ventricle; IV: intravenous injection. **B.** Survival curves. 2 experiments pooled; n= 36 larvae for CTRL and OV, 31 for IV, and 33 for HBV injection. **C**) bacterial burden evaluated over time by fluorescent pixel counts. 1 experiment, 6 larvae per condition. **D.** Representative images of *L. pneumophila* dissemination, determined by live imaging using a fluorescence stereomicroscope, of zebrafish larvae infected with a HD WT-, in closed compartments (OV, HBV) or in the bloodstream (IV). Infected larvae were live imaged 4h, 24h, 48h, and 72h post *L. pneumophila* injection. Only GFP fluorescence is shown. Green autofluorescence of the lens eye (e) or of the gastrointestinal tract (g) is indicated on CTRL larvae. $P < 0.05$ was considered statistically significant (symbols: **** $P < 0.0001$; ***$P < 0.001$; **$P < 0.01$; *$P < 0.05$). No symbol on graphs means that not statistically differences were observed.

**Figure S4: Bath immersion of 120 hpf zebrafish larvae using *WT L pneumophila* infected *A castellanii***
**A)** Survival curves. **B)** % of larvae with GFP bacteria. A and B: 1 experiment, 30 larvae for WT Lpp-amoebae, 10 for WT Lpp and 10 for amoebae. **C)** Representative fluorescent imaging of larvae with GFP bacteria in the intestinal tract followed over time. The intestinal tractus is highlighted with white dotted lines. Arrowhead points to GFP bacteria being eliminated with the fecal content. D) representative closeup of GFP bacteria in the intestinal tract. $P < 0.05$ was considered statistically significant (symbols: **** $P < 0.0001$; ***$P < 0.001$; **$P < 0.01$; *$P < 0.05$). No symbol on graphs means that not statistically differences were observed.

**Figure S5: *Δspl* mutant injected in the yolk and *L. pneumophila* WT but not the T4SS mutant proliferates in the yolk of zebrafish and the yolk of chicken eggs upon direct injection. A).** 72hpf larva: the yolk cell is highlighted in blue and the yolk content in pink**. B).** Survival curves of 72hpf larvae upon injection in the yolk of HD WT (red curve), Δ*dot*A (green curve) or *spi* mutant (violet curve) *L. pneumophila* strain. CTRL larvae (black curve). One experiment, 24 larvae for each condition. Significant differences are indicated with stars. **C)** Survival curves of embryonated chicken eggs inoculated with WT strain (in red, n=9), Δ*dot*A strain (in blue, n=7) or PBS (in grey, n=7). Survival expressed in percentage and time in days. Comparison of survival curves was performed using Log-rank (Mantel-Cox) test. $P < 0.05$ was considered statistically significant. **D)** Quantification of *L. pneumophila*, expressed in $\log_{10}$ CFU/mL, in yolk sac of WT-infected embryos (n=9) and Δ*dotA*-infected embryos (n=6). Dead embryos are represented with black dots and those euthanized (alive at day 6) with white dots. The inoculum in the yolk sac after infection was estimated by taking into account the count of *L. pneumophila* in the inoculum (WT and Δ*dotA*) before injection and the volume of the yolk sac. Comparison of *L. pneumophila* quantifications between WT- and Δ*dotA*-infected embryos was done using Mann-Whitney test. $P < 0.05$ was considered statistically significant (symbols: **** $P < 0.0001$; ***$P < 0.001$; **$P < 0.01$; *$P < 0.05$).

**Figure S6: Quantification of bacterial burden in the whole body, in the body or in the yolk region versus macrophage or neutrophil quantification in the body or in the yolk region in HD WT-GFP infected larvae followed overtime**. Two independent experiments plotted for each phagocyte type (total of 11 larvae for macrophage or 11 larvae for neutrophil quantification). Quantification of the fluorescent images (GFP bacteria and RFP leukocytes) was done using CellProfiler software (see Material and Methods for details about the pipeline). Bacterial burden quantification was done over the whole larva (red dot) or discriminating the body (light blue dot) form the yolk region (pink dot). Scheme of 72hpf with body (light blue) and yolk region (pink) highlighted; the yolk sustaining *L. pneumophila* growing has been indicated with green dots**.** Quantification of macrophage or neutrophil located in body (light blue dot) or yolk (pink dot) over time. $P < 0.05$ was considered statistically significant (symbols: **** $P < 0.0001$; ***$P < 0.001$; **$P < 0.01$; *$P < 0.05$). No symbol on graphs means that not statistically differences were observed.

**Figure S7: Macrophage and neutrophil depletion by morpholino: evaluation of the impact on the non-depleted leukocyte population**. Comparison of the impact of *spi1b* morpholino injection that blocks macrophage development or csf3r morpholino injection that blocks neutrophil development were administered. Macrophages (red symbols) and neutrophils (green symbols) were counted in CTRL (open symbols) or morphant (full symbols) conditions. **A)** effect of spe1b morpholino on macrophages and neutrophils, showing that spe1b morpholino injection leads to the specific depletion of macrophages and not neutrophils. Related to Fig 4: 2 plotted experiments, n=10 larvae per group. B) effect of Csf3R morpholino on macrophages and neutrophils, showing that Csf3R morpholino injection leads to the specific depletion neutrophils and slightly impairs the number of macrophages. Related to Fig 5: 2 plotted experiments, n=10 larvae per group

**Figure S8: A. Correlation between bacterial burden (evaluated by fluorescence on individual injected larvae before RNA extraction) and cytokine gene induction at 48 and 72hpi upon bloodstream injection LD, HD WT or HD Δ*dot*A *L pneumophila* strain.** Control non injected, HD WT or HD Δ*dot*A injected larvae were scored under the fluorescent microscope for evaluating bacterial burden immediately before to be lysed and processed for RNA extraction. "-", "+" to "+++ " respectively indicate no or in, creasing bacterial burden. "-" and "+" symbols were also used to respectively indicate infected dead or live larvae. Related to Fig 7A-D. **C-D) Cytokine gene (*il1b*, *tnfa, ifng1/2* ) induction is independent from Myd88 signalling in *L pneumophila* HD WT infected zebrafish larvae** .Cytokine gene induction was measured from individual *myd88*$^{hu3568}$ mutant larvae injected with a HD (red curves) of WT-GFP and non-injected fish (CTRL, black curves). The same colours are used for individual CTRL non injected (black dashed) or HD WT injected (red dashed) zebrafish curves. Data plotted are from one experiment (n=5 larvae for each condition); individual values are shown, and curves correspond to the medians. There is no statistically significant difference between CTRL and *myd88*$^{hu3568}$ mutant curves over time for all the conditions analysed. Related to Fig 7 E,F.

**Movies are available at** https://zenodo.org/record/7307933

**Legends:**

**Movie S1: *L. pneumophila* growing in the yolk region at 72hpi: localization in the yolk in AB wild type larva.** AB wild type larva 72hpf was injected in the bloodstream with HD of *L. pneumophila* WT-GFP, and was analyzed using confocal high microscopy at 72hpi, to study the behavior of the highly growing bacteria in the yolk region. The infected larva was mounted laterally and acquired using a 20X oil-immersion objective. The acquired Z-stack was deconvolved using Leica Lightening Plug-in and processed for 3D visualization and volume rendering, using IMARIS 9.6 (Bitplane). Note the complex, filamentous, highly aggregate structures (green) formed by the growing *Legionella* in the yolk region (visualized by the bright field).

**Movie S2: *L. pneumophila* growing in the yolk region at 72hpi: interactions with blood vessels.** *kdrl*:mCherry (red blood vessels) 72hpf larva was injected in the bloodstream with HD of *L. pneumophila* WT-GFP, and was analyzed with confocal high microscopy at 72hpi, to study the behavior of the highly growing bacteria in the yolk region and their interactions with the yolk vasculature. The infected larva was mounted laterally and acquired using a 20X oil-immersion objective. The acquired Z-stack was deconvolved using Leica Lightening Plug-in and processed for 3D visualization and volume rendering, using IMARIS 9.6 (Bitplane). The interactions of the blood vessels (red cells) with the growing bacterial aggregates (green), and the yolk region (bright field) are shown at various magnifications (scale bar indicated on the movie) and various rotations angles to highlight the complex filamentous bacterial structures and their interactions with the blood vessels. Due to the peculiar yolk composition and thickness, it was impossible to acquire the fluorescence of the bacteria growing inside the yolk region distal to the objective, thus appearing as big dark spots.

**Movie S3: *L. pneumophila* growing in the yolk region at 72hpi: interactions with macrophages** *Mfap4*: mCherry (red macrophages) 72hpf larva was injected in the bloodstream with HD of *L. pneumophila* WT-GFP, and was analyzed with confocal high microscopy at 72hpi, to study the behavior of the bacteria growing in the yolk region and their interactions with macrophages. The infected larva was mounted ventrally and acquired using a 40X water-immersion objective. Only the yolk region containing the bacterial aggregates was imaged. The acquired Z-stack was deconvolved using Leica Lightening Plug-in and processed for 3D visualization and volume rendering, using IMARIS 9.6 (Bitplane). The interactions of macrophages (red cells) with the growing bacterial aggregates (green), and the yolk region (bright field) are shown at various magnifications (scale bar indicated on the movie) and various rotation angles to highlight the complex filamentous bacterial structures and the recruited macrophages, that recognize the growing bacteria, but fail to penetrate the yolk content, and to engulf the bacterial aggregates. Due to the peculiar yolk composition and thickness, it was impossible to acquire the fluorescence of the bacteria growing inside the yolk distal to the objective, thus appearing as big dark spots.

**Movie S4: *L. pneumophila* growing in the yolk region at 72hpi: interactions with neutrophils.** *Lys*:DsRed (red neutrophils) 72hpf larva was injected in the bloodstream with HD of *L. pneumophila* WT-GFP*,* and was analyzed with confocal high microscopy at 72hpi, to study the behavior of the bacteria growing in the yolk region and their interactions with neutrophils. The infected larva was mounted laterally and acquired using a 20X oil-immersion objective. The acquired Z-stack was deconvolved using Leica Lightening Plug-in and processed for 3D visualization and volume rendering, using IMARIS 9.6 (Bitplane). The interactions of neutrophils (red cells) with the growing bacterial aggregates (green), and the yolk region (bright field) are shown at various magnifications (scale bar indicated on the movie) and various rotations angles to highlight the complex filamentous bacterial structures and the recruited neutrophils, that recognize and sense the growing bacteria, migrate to them, but fail to penetrate the yolk content, and to engulf the big bacterial aggregates. Due to the peculiar yolk composition and thickness, it was impossible to acquire the fluorescence of the bacteria growing inside the yolk distal to the objective, thus appearing as big dark spots**.**

**Movie S5:** *L. pneumophila* growing in the yolk region between 48 and 72hpi: interactions with macrophages (related to Figure 3E). 4D, 40X objective, 6h time lapse between 48-72hpi, 1microm optical sections, infected larva mounted ventral, bacteria growing in aggregate on the yolk. *Mfap4*: mCherry (red macrophages) 72hpf larva was injected in the bloodstream with HD of *L. pneumophila* WT-GFP*,* and was analyzed with confocal high microscopy between 48 and 72hpi , to study the behavior of the bacteria growing in the yolk region, their spatial localization (above or below the yolk plasma membrane), and their interactions with macrophages overtime. The infected larva was mounted ventrally and acquired using a 40X water-immersion objective. Only the yolk region containing the bacterial aggregates was imaged. The acquired Z-stack was deconvolved using Leica

Lightening Plug-in and processed for 4D visualization and volume rendering, using IMARIS 9.6 (Bitplane). Macrophage and bacteria surfacing automatically done, yolk region manually delimitated frame by frame with Imaris.

**Movie S6: Macrophage - *L. pneumophila* interactions (LD, HD).** *Mfap4*: mCherry (red macrophages) 72hpf larvae were injected in the bloodstream with LD (left panel) or HD (middle panel) of *L. pneumophila* WT-GFP or with HD OF *L. pneumophila* Δ*dotA*-GFP (right panel), mounted laterally and acquired using high resolution confocal microscopy to analyze macrophages (red cells) bacteria (green) interactions immediately upon bacteria injection. The infected larvae were acquired over time from 20 min to approximately 16 hours post injection. Maximum projections of the acquired Z-stacks are shown. The 3D movies generated were combined using Image J software, to have them side by side, to compare the macrophage-bacteria interaction over time in the various conditions. Left panel: *mfap4*: mCherry (red macrophages) 72hpf larva injected in the bloodstream with LD wt GFP *Legionella* (green). Note that macrophages are recruited to the injected bacteria, engulf them, and the bacteria are cleared progressively from the bloodstream. Middle panel: *mfap4*: mCherry (red macrophages) 72hpf larva injected in the bloodstream with HD wt GFP *Legionella* (green). Macrophages are recruited upon bacteria injection but failed to eliminate them over time; the phagocytosing macrophages round-up. Right panel: *mfap4*: mCherry (red macrophages) 72hpf larva injected in the bloodstream with HD GFP Δ*dotA Legionella* (green). Note that the recruited macrophages efficiently engulf and eliminate the injected bacteria, clearing them progressively from the blood and the mesenchyme near the point of injection.

**Movie S7: Neutrophil - *L. pneumophila* (LD, HD) interactions**. (*Lys*: DsRed (red neutrophils) 72hpf larvae were injected in the bloodstream with LD (left panel) or HD (middle panel) *L. pneumophila* WT-GFP, or with HD of Δ*dotA*-GFP (right panel), mounted laterally and acquired using high resolution confocal microscopy to analyze neutrophil (red cells) bacteria (green) interactions immediately upon bacteria injection. The infected larvae were acquired over time from 20 min to approximately 16 hours post injection. Maximum projections of the acquired Z-stacks are shown. The 3D movies generated were combined using ImageJ software, to have them side by side, to compare neutrophil-bacteria interactions over time in the various conditions. Left panel: *Lys*: DsRed (red neutrophils) 72hpf larva injected in the bloodstream with LD of WT-GFP (green). Note that neutrophils are recruited to the injected bacteria, engulfing the bacteria trapped in the mesenchyme near the site of injection, cooperating with macrophages (DsRed – cells, GFP+ having engulfed large amount of GFP bacteria), clearing progressively the infection. Middle panel: *Lys*:DsRed (red neutrophils) 72hpf larva injected in the bloodstream with HD of WT-GFP (green). Neutrophils are massively recruited upon bacterial injection but failed to eliminate them over time; the phagocytosing neutrophils round-up and loose DsRed fluorescence, suggesting cell death. Right panel: *lys:*DsRed (red neutrophils) 72hpf larva injected in the bloodstream with HD of Δ*dotA*-GFP *L. pneumophila* (green). Note that the recruited neutrophils engulf and eliminate the injected bacteria, clearing them progressively from mesenchyme near the point of injection, efficiently cooperating with macrophages in controlling the infection**.**

**Movie S8: Dying phagocytosing neutrophils upon *L. pneumophila* HD injection (related to Figure 5).** (*Lys*: DsRed (red neutrophils) 72hpf larvae were injected in the bloodstream with HD *L. pneumophila* WT-GFP, mounted laterally and acquired using high resolution confocal microscopy to analyze neutrophil (red cells) bacteria (green) interactions immediately upon bacteria injection. The infected larvae were acquired over time from 20 min to approximately 16 hours post injection. Time lapses every 1'30". Maximum projections of the acquired Z-stacks (2mm per optical section) are shown. 6 neutrophils were manually tracked with Fiji (1 to 6) and highlighted with open white circle overtime. Note that the tracked neutrophils having engulfed *L. pneumophila* progressively dyed, rounding up and losing their red fluorescence, while the green fluorescence of the GFP bacteria is still visible overtime.

**Figure S1**

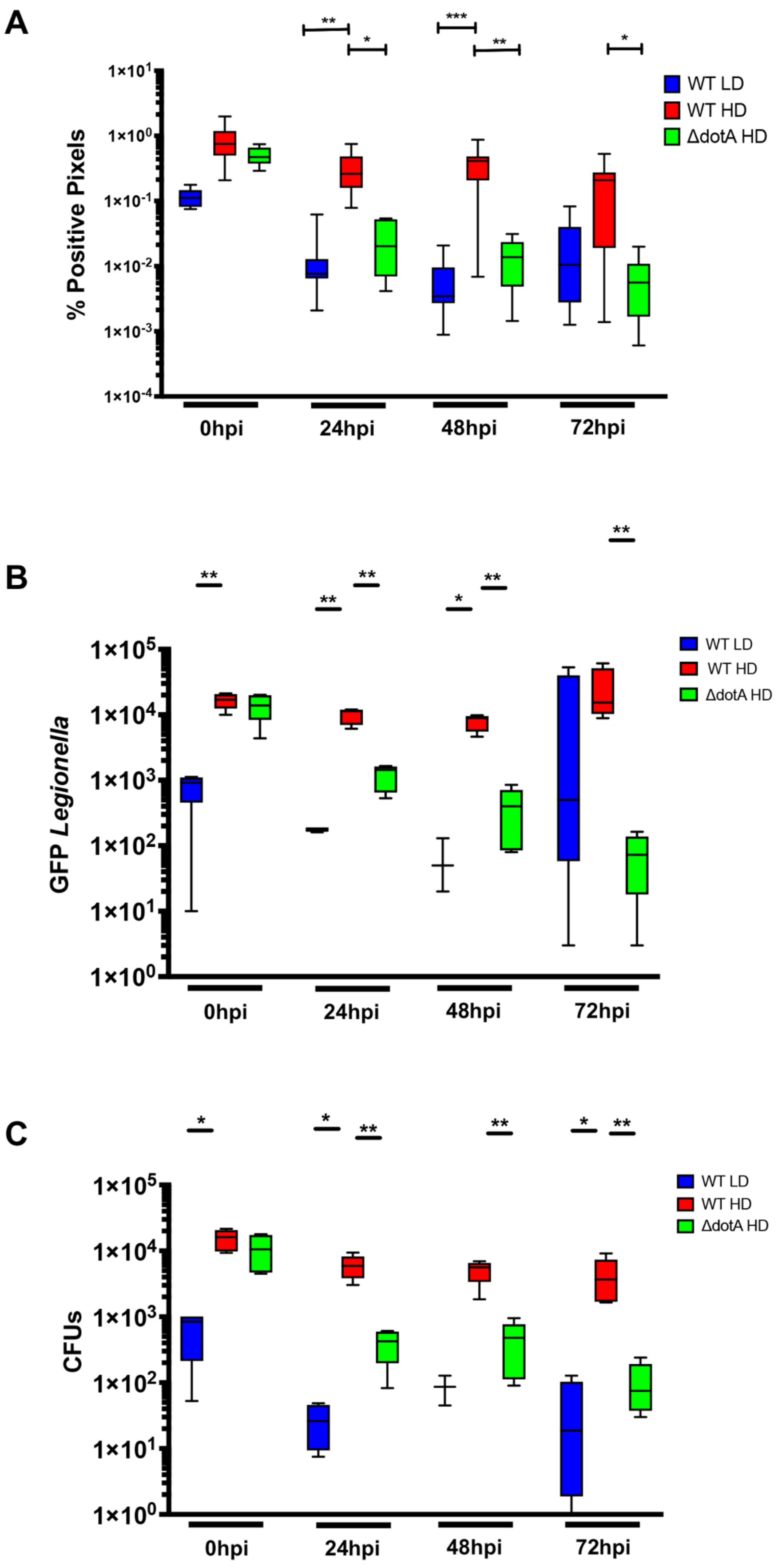
A
% Positive Pixels
1×10^1
1×10^0
1×10^-1
1×10^-2
1×10^-3
1×10^-4
WT LD
WT HD
ΔdotA HD
0hpi
24hpi
48hpi
72hpi
B
GFP Legionella
1×10^5
1×10^4
1×10^3
1×10^2
1×10^1
1×10^0
WT LD
WT HD
ΔdotA HD
0hpi
24hpi
48hpi
72hpi
C
CFUs
1×10^5
1×10^4
1×10^3
1×10^2
1×10^1
1×10^0
WT LD
WT HD
ΔdotA HD
0hpi
24hpi
48hpi
72hpi

**Figure S2**

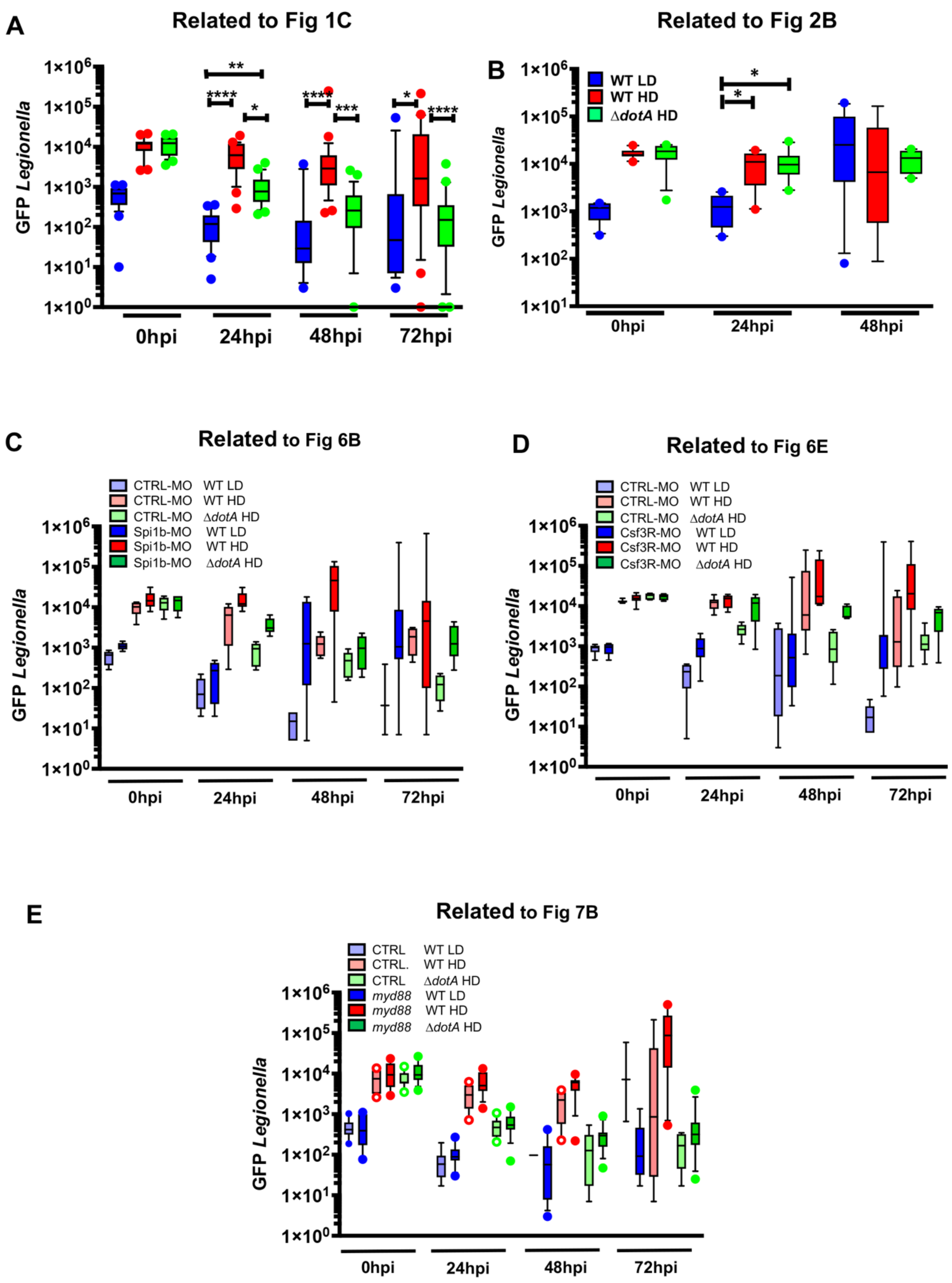
A
Related to Fig 1C
B
Related to Fig 2B
WT LD
WT HD
ΔdotA HD
GFP Legionella
0hpi
24hpi
48hpi
72hpi
C
Related to Fig 6B
CTRL-MO WT LD
CTRL-MO WT HD
CTRL-MO ΔdotA HD
Spi1b-MO WT LD
Spi1b-MO WT HD
Spi1b-MO ΔdotA HD
D
Related to Fig 6E
CTRL-MO WT LD
CTRL-MO WT HD
CTRL-MO ΔdotA HD
Csf3R-MO WT LD
Csf3R-MO WT HD
Csf3R-MO ΔdotA HD
E
Related to Fig 7B
CTRL WT LD
CTRL. WT HD
CTRL ΔdotA HD
myd88 WT LD
myd88 WT HD
myd88 ΔdotA HD

**Figure S3**

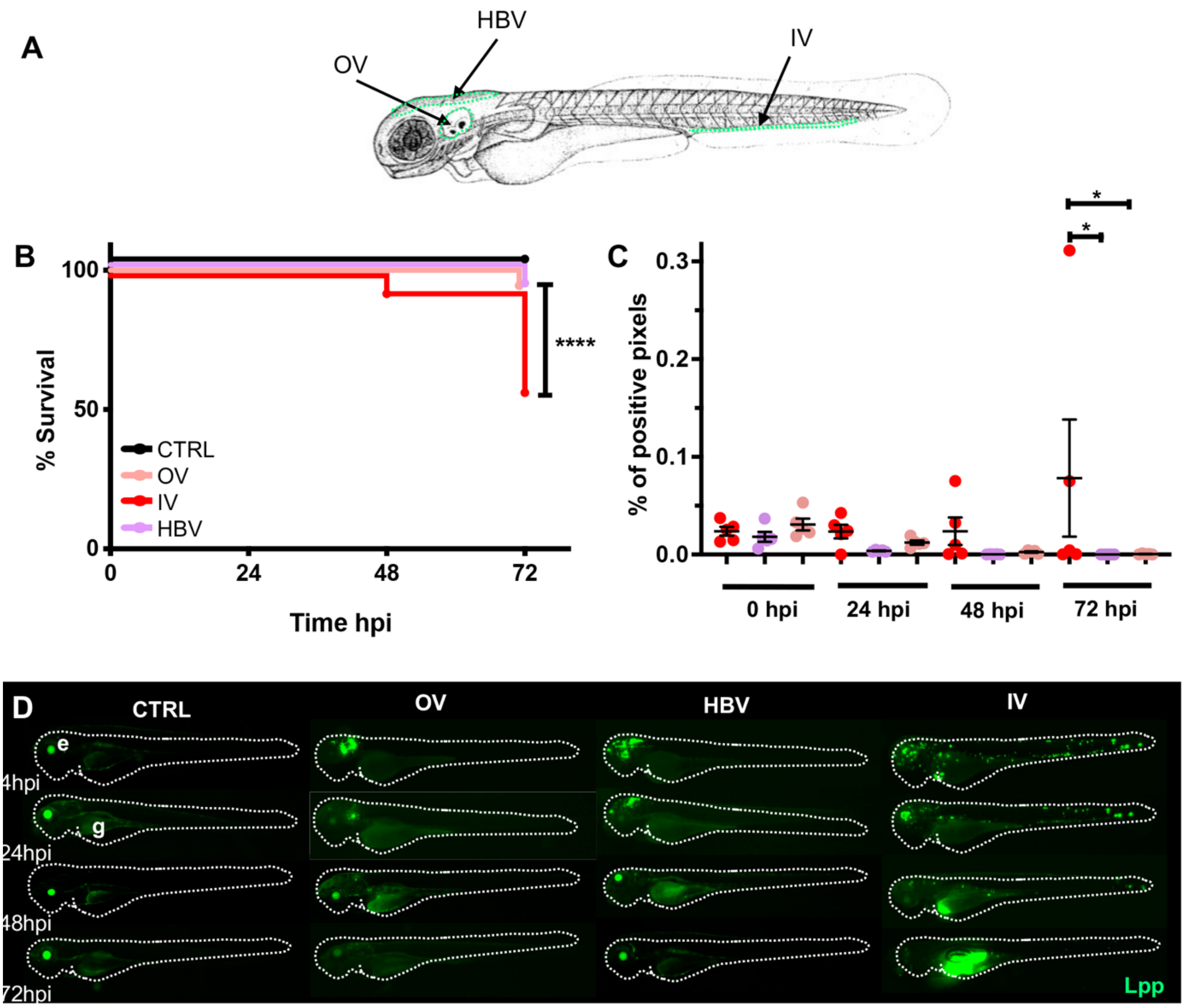
A
HBV
OV
IV
B
100
50
0
% Survival
0
24
48
72
Time hpi
****
CTRL
OV
IV
HBV
C
0.3
0.2
0.1
0.0
% of positive pixels
*
*
0 hpi
24 hpi
48 hpi
72 hpi
D
CTRL
OV
HBV
IV
e
g
4hpi
24hpi
48hpi
72hpi
Lpp

**Figure S4**

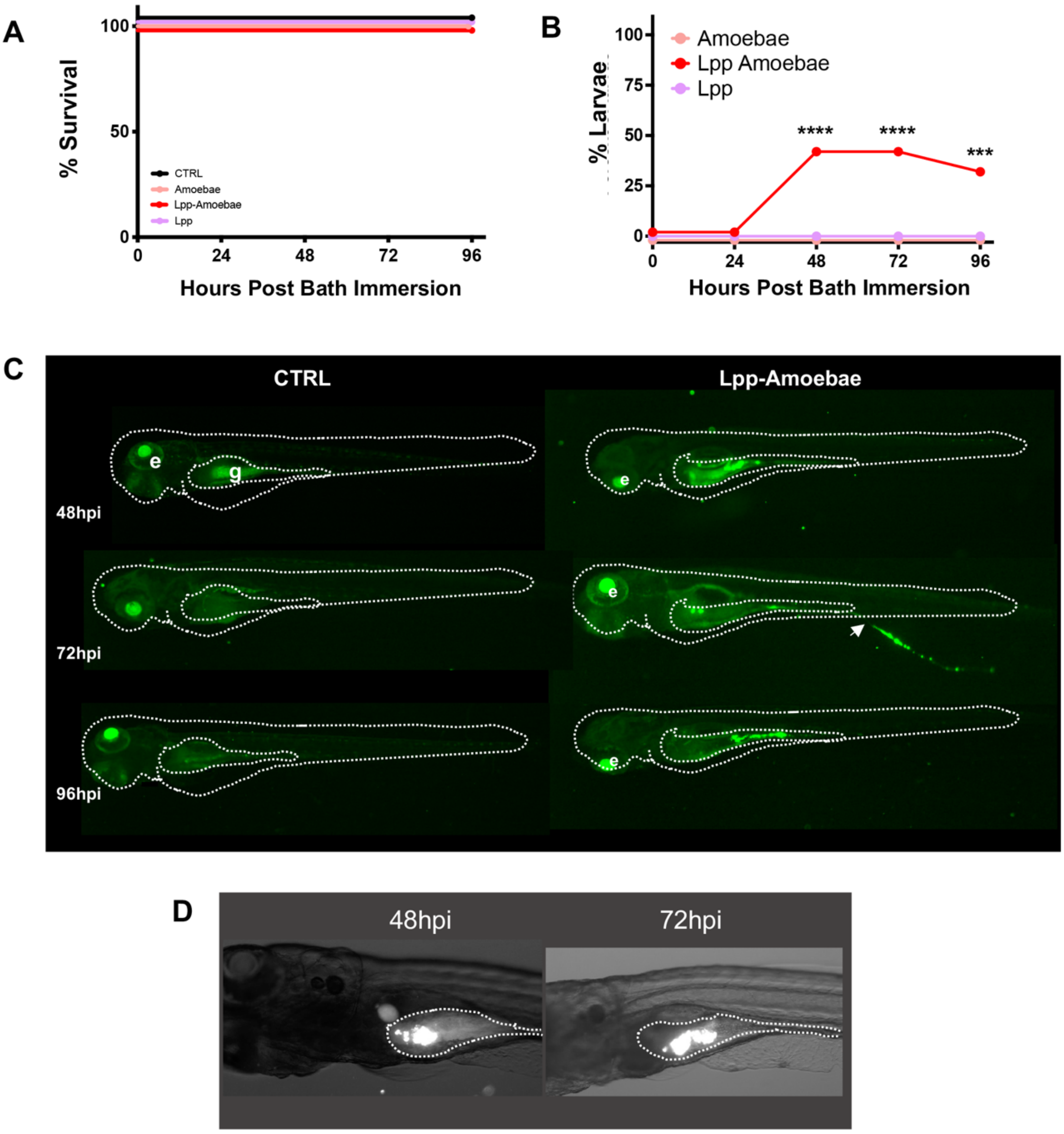
A
100
50
0
% Survival
CTRL
Amoebae
Lpp-Amoebae
Lpp
0
24
48
72
96
Hours Post Bath Immersion
B
100
75
50
25
0
% Larvae
Amoebae
Lpp Amoebae
Lpp
****
****
***
Hours Post Bath Immersion
C
CTRL
Lpp-Amoebae
e
g
48hpi
72hpi
96hpi
D
48hpi
72hpi

**Figure S5**

A

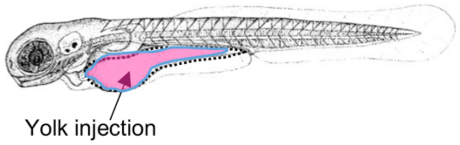


B

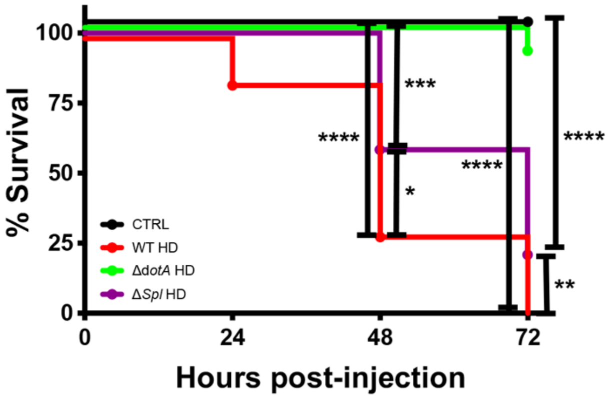


C

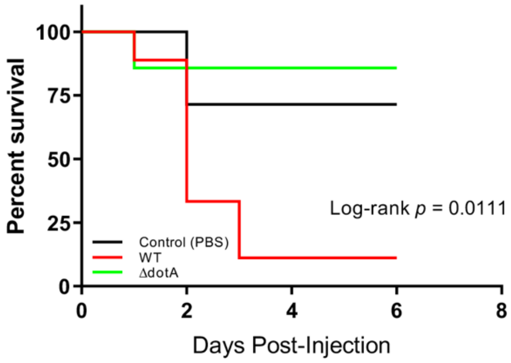


D

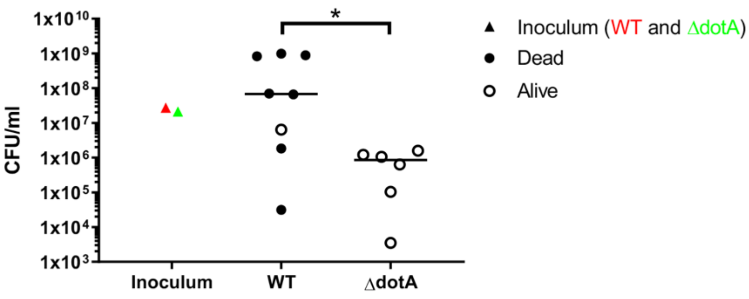

**Figure S6**

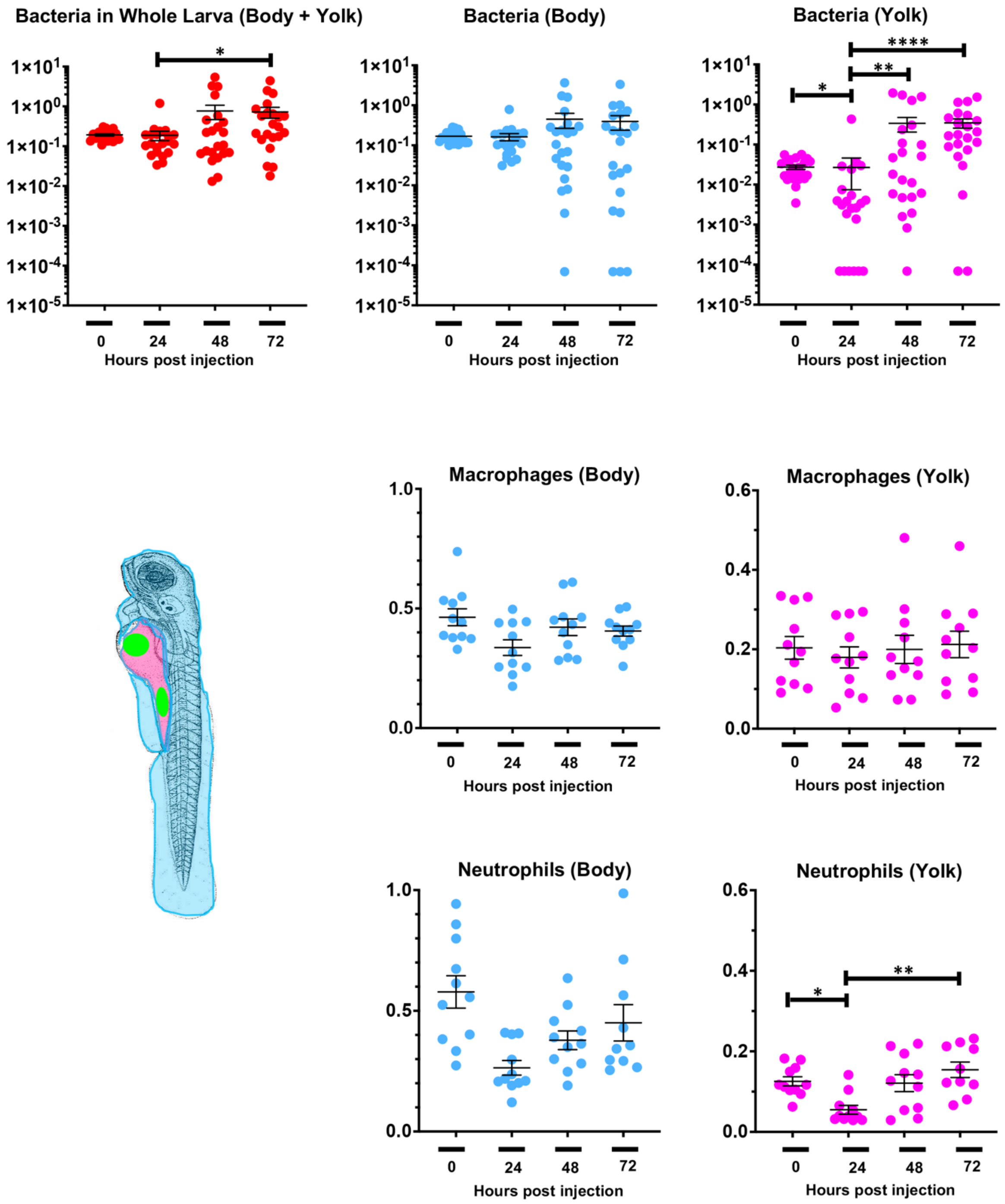
Bacteria in Whole Larva (Body + Yolk)
Bacteria (Body)
Bacteria (Yolk)
Macrophages (Body)
Macrophages (Yolk)
Neutrophils (Body)
Neutrophils (Yolk)
Hours post injection
0
24
48
72

**Figure S7**

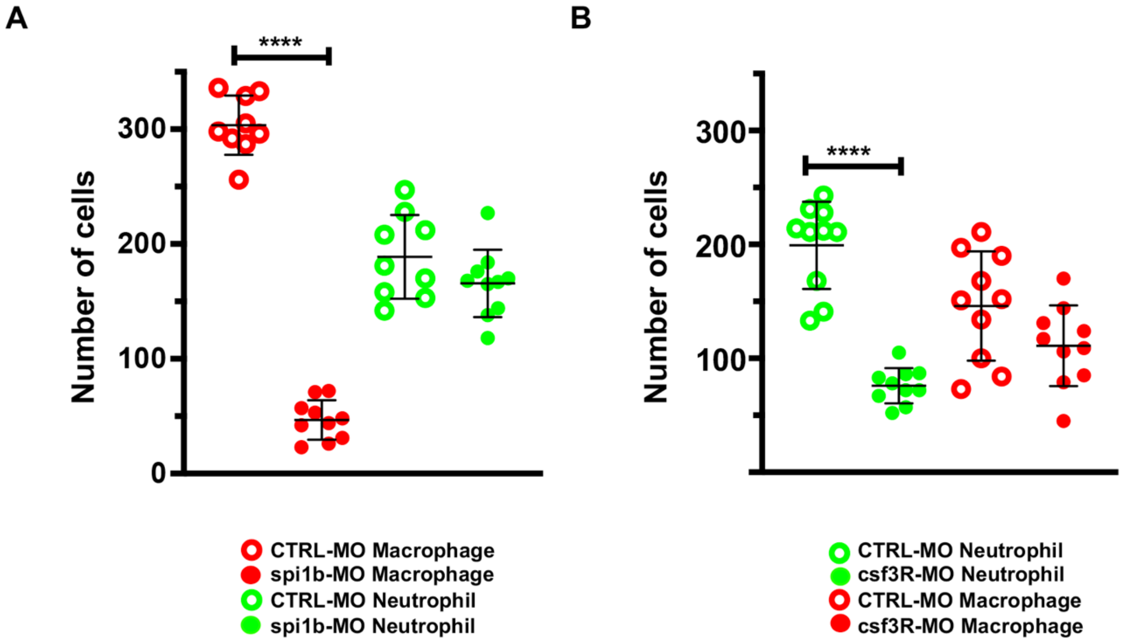
A
****
Number of cells
300
200
100
0
CTRL-MO Macrophage
spi1b-MO Macrophage
CTRL-MO Neutrophil
spi1b-MO Neutrophil
B
****
Number of cells
300
200
100
CTRL-MO Neutrophil
csf3R-MO Neutrophil
CTRL-MO Macrophage
csf3R-MO Macrophage

**Figure S8**

**A**

| 48hpi | Live | Lpp | il1b | tnfa | ifng1 |
|---|---|---|---|---|---|
| CTRL 1 | + | - | | | |
| CTRL 2 | + | - | 0,25 | 0,13 | 0,75 |
| CTRL 3 | + | - | 0,26 | 0,30 | 0,46 |
| CTRL 4 | + | - | 0,34 | 0,28 | 1,45 |
| CTRL 5 | + | - | 0,59 | 0,67 | 0,87 |
| WT LD 1 | + | + | 0,30 | 2,62 | 2,09 |
| WT LD 2 | + | + | 0,74 | 2,52 | 4,31 |
| WT LD 3 | + | + | | | |
| WT LD 4 | + | + | 0,34 | 1,44 | 1,29 |
| WT LD 5 | + | - | 0,91 | 0,92 | 0,22 |
| WT HD 1 | + | + | 0,96 | 4,96 | 0,13 |
| WT HD 2 | + | ++ | 9,32 | 3,47 | 1,48 |
| WT HD 3 | +/- | +++ | 44,36 | 2,68 | 28,15 |
| WT HD 4 | + | ++ | 6,57 | 3,72 | 1,08 |
| WT HD 5 | - | +++ | 219,66 | 2651,60 | 1676,55 |
| ΔdotA HD 1 | + | + | 2,80 | 2,19 | 0,94 |
| ΔdotA HD 2 | + | - | 2,03 | 1,09 | 2,25 |
| ΔdotA HD 3 | + | + | 3,54 | 2,16 | 0,71 |
| ΔdotA HD 4 | + | - | 3,05 | 2,58 | 1,58 |
| ΔdotA HD 5 | + | - | 3,50 | 3,75 | 1,95 |

| 72hpi | Live | Lpp | il1b | tnfa | ifng1 |
|---|---|---|---|---|---|
| CTRL 1 | + | - | 0,16 | 0,19 | 0,21 |
| CTRL 2 | + | - | 0,40 | 0,35 | 0,54 |
| CTRL 3 | + | - | 0,24 | 0,52 | 0,41 |
| CTRL 4 | + | - | 0,20 | 0,04 | 0,72 |
| CTRL 5 | + | - | | | |
| WT LD 1 | + | - | 0,22 | 0,44 | 0,20 |
| WT LD 2 | + | + | 0,25 | 0,35 | 0,64 |
| WT LD 3 | + | - | 0,67 | 0,61 | 1,14 |
| WT LD 4 | + | + | 0,47 | 0,32 | 2,49 |
| WT LD 5 | + | - | 0,50 | 0,47 | 4,52 |
| WT HD 1 | + | +++ | 631,15 | 907,63 | 2386,75 |
| WT HD 2 | - | +++ | 84,48 | 3269,18 | 1146,69 |
| WT HD 3 | -- | ++ | 15,10 | 781,61 | 623,30 |
| WT HD 4 | - | +++ | 166,38 | 9894,70 | |
| WT HD 5 | - | ++ | 101,58 | 83,90 | 0,69 |
| ΔdotA HD 1 | + | + | | | |
| ΔdotA HD 2 | + | - | | | |
| ΔdotA HD 3 | + | + | 0,75 | 1,26 | 0,85 |
| ΔdotA HD 4 | + | - | 0,54 | 1,14 | 0,77 |
| ΔdotA HD 5 | + | - | 1,87 | 2,56 | 1,81 |

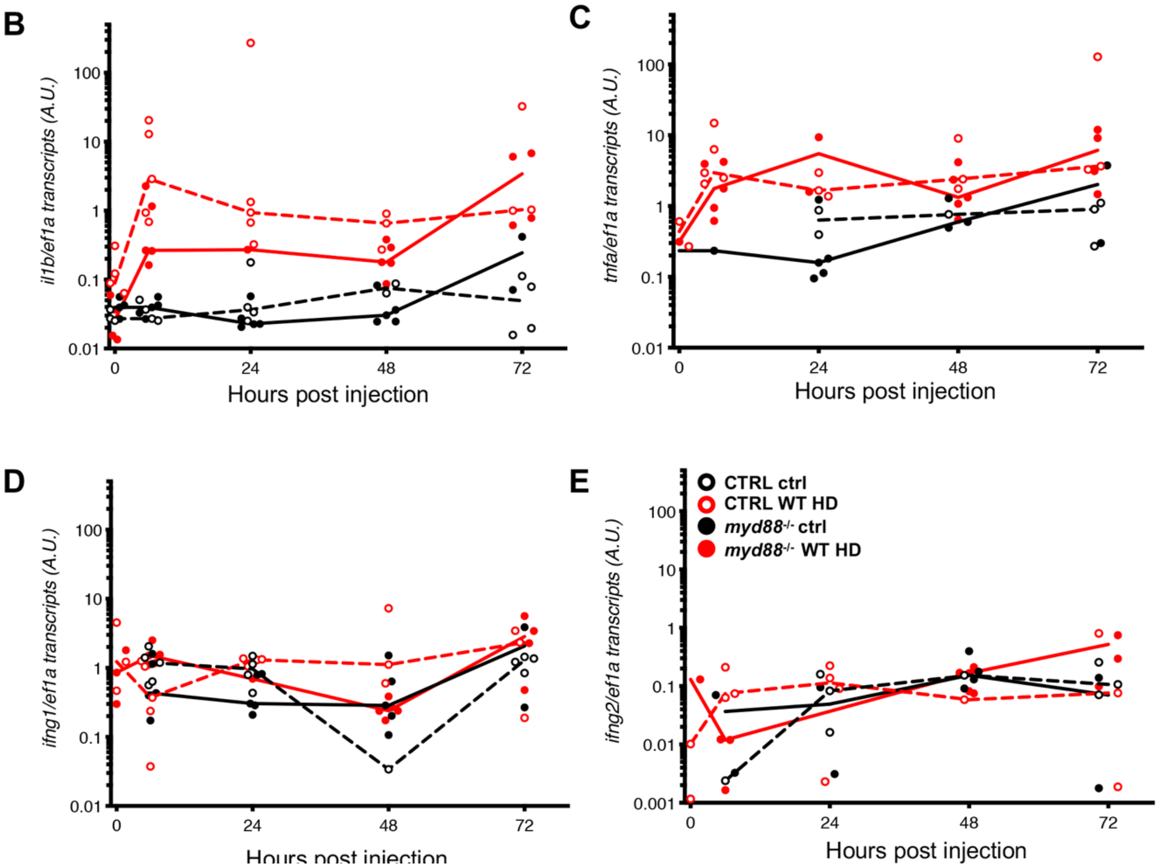
